# Agtrevirus phage AV101 infect diverse extended spectrum β-lactamase *E. coli* by recognizing four different O-antigens

**DOI:** 10.1101/2023.09.19.558411

**Authors:** Anders Nørgaard Sørensen, Dorottya Kalmar, Veronika Theresa Lutz, Victor Klein-Sousa, Nicholas M. I. Taylor, Martine C. Holst Sørensen, Lone Brøndsted

**Affiliations:** Department of Veterinary and Animal Sciences, University of Copenhagen, Frederiksberg C, Denmark; Structural Biology of Molecular Machines Group, Protein Structure & Function Program, Novo Nordisk Foundation Center for Protein Research, Faculty of Health and Medical Sciences, University of Copenhagen, Copenhagen, Denmark

**Keywords:** Bacteriophages, *Ackermannviridae*, *Agtrevirus*, host range analysis, tail spike proteins, O-antigen receptors

## Abstract

Bacteriophages in the *Agtrevirus* genus are known for expressing multiple tail spike proteins (TSPs), but little is known about their genetic diversity and host recognition apart from their ability to infect diverse *Enterobacteriaceae* species. Here we aim to determine the genetic differences that may account for the diverse host ranges of *Agrevirus* phages. We performed comparative genomics of 14 *Agtrevirus* and identified only a few genetic differences including genes involved in nucleotide metabolism. Most notably was the diversity of the *tsp* gene cluster, specifically in the receptor binding domains that were unique among most of the phages. We further characterized agtrevirus AV101 infecting nine diverse Extended Spectrum β-lactamase (ESBL) *E. coli* and demonstrated that this phage encoded four unique TSPs among *Agtrevirus*. Purified TSPs formed translucent zones and inhibited AV101 infection of specific hosts, demonstrating that TSP1, TSP2, TSP3, and TSP4 recognize O8, O82, O153, and O159 O-antigens of ESBL *E. coli*, respectively. BLASTp analysis showed that the receptor binding domain of TSP1, TSP2, TSP3 and TSP4 are similar to TSPs encoded by *E. coli* prophages and distant related virulent phages. Thus, *Agtrevirus* may have gained their receptor binding domains by recombining with prophages or virulent phages. Overall, combining bioinformatic and biological data expands the understanding of TSP host recognition of *Agtrevirus* and give new insight into the origin and acquisition of receptor binding domains of *Ackermannviridae* phages.

**One sentence summary:** Agtrevirus phage AV101 express four unique tail spike proteins that recognize different O-antigens of Extended Spectrum β-Lactamase producing *E. coli*.

## Introduction

The recent establishment of genome-based taxonomy of phages has led to the classification of an increasing number of phage families and genera (Lefkowitz *et al*. 2018). Thus, based on genetic similarity, the *Ackermannviridae* family was established in 2017 and currently includes two subfamilies (*Cvivirinae* and *Aglimvirinae)* and ten genera (*Kuttervirus*, *Agtrevirus, Limestonevirus Taipeivirus*, *Tedavirus*, *Nezavisimistyvirus*, *Miltonvirus*, *Campanilevirus*, *Vapseptimavirus* and *Kujavirus)* (Kropinski *et al*. 2017; Adriaenssens *et al*. 2018). In the *Ackermannviridae* family, phages exhibit a conserved genome architecture including gene synteny in genomes of substantial size of approximately 150 kilobases. In addition, these phages encode hydroxymethyluracil (HMdU) synthase leading to substitution of thymine with hydroxylmethyl uracil in their genomes (Kutter *et al*. 2011; Adriaenssens *et al*. 2012b, 2012a; Hsu *et al*. 2013). This nucleotide substitution has been suggested to prevent cleavage of the phage genomes by host-encoded restriction enzymes, which has been proposed to allow *Ackermannviridae* phages to infect a broad range of *Enterobacteriaceae* species (Kutter *et al*. 2011; Adriaenssens *et al*. 2012b, 2012a; Hsu *et al*. 2013).

The most well-studied feature of *Ackermannviridae* phages is their receptor binding properties arising from expressing up to four diverse tail spike proteins (TSPs). These four TSPs form a complex protruding from the baseplate in a star-like morphology that can be visually observed in transmission electron micrographs (TEMs) (Adriaenssens *et al*. 2012a; Plattner *et al*. 2019). The TSPs are hinged together in a complex, and to the baseplate, by protein interactions between the conserved N-terminal modules of each TSPs (Plattner *et al*. 2019; Chao *et al*. 2022). In contrast, the receptor-binding domain of these TSPs are highly diverse and binds to polysaccharide receptors like the O-antigen or K-antigen through conserved folds including a β-helix commonly observed for TSPs in the *Caudoviricetes* order (Barbirz *et al*. 2008; Andres *et al*. 2010; Lee *et al*. 2017; Olszak *et al*. 2017; Prokhorov *et al*. 2017; Kunstmann *et al*. 2018). As the phages express multiple TSPs, each capable of binding to a specific O-antigen or K-antigen, their host ranges are broader compared to phages only expressing a single TSP targeting such polysaccharide receptor (Steinbacher *et al*. 1996; Plattner *et al*. 2019; Sørensen *et al*. 2021; Witte *et al*. 2021). For example, *Kuttervirus* CBA120 infects *Salmonella* O21*, E. coli* O157, *E. coli* O77 and *E. coli* O78 strains through the specific binding of TSP1, TSP2, TSP3 and TSP4, respectively (Plattner *et al*. 2019). Both *E. coli* and *Salmonella* are common hosts for *Kuttervirus* and *Agtrevirus* phages, but these bacteria express more than 185 and 46 diverse O-antigens, respectively (Liu *et al*. 2014, 2020). Thus, *Ackermannviridae* phages match the diversity of O-antigens expressed the bacterial hosts by encoding diverse TSP. For example, we previously analysed 374 TSPs encoded by 99 *Ackermannviridae* phages and found 96 diverse TSP subtypes each carrying unique receptor binding domains (Sørensen *et al*. 2021). Furthermore, the TSP subtypes were strongly associated with phage genera. In addition, further analysis allowed us to predict the host recognition of TSP subtypes encoded by phages of the *Kuttervirus, Limestonevirus* and *Taipei* genera, but due to lack of biological data, no information of *Agtrevirus* phages could be revealed (Sørensen *et al*. 2021).

The *Agtrevirus* genus is not very well studied in terms of host range and receptor recognition. The phages are known to infect Gram-negative bacteria like *Salmonella*, *E. coli*, *Enterobacter* and *Shigella*. But, unlike other members of the *Ackermannviridae* family, such as phages in *Kuttervirus* genus that infect both *E. coli* and *Salmonella*, *Agtrevirus* phages are only known to infect one bacterial species each (Anany *et al*. 2011; Heyse *et al*. 2015; Soffer *et al*. 2016; Akter *et al*. 2019; Thanh *et al*. 2020; Kwon *et al*. 2021; Imklin *et al*. 2022). We recently isolated agtrevirus AV101 infecting Extended Spectrum β-Lactamase (ESBL) *E. coli* (Vitt *et al*. 2023). A large host range analysis of 198 ESBL *E. coli* strains showed that *Agtrevirus* phage AV101 had a narrow host range only infecting 9 of 198 strains tested (Vitt *et al*. 2023). In addition, we previously observed that five *Agtrevirus* phages express unique TSPs without similarity to other phages in the genus or *Ackermannviridae* family (Sørensen *et al*. 2021). Thus, the diverse hosts infected by *Agtrevirus* phages may be due to genetic differences between the phages including unique receptor binding domains of the TSPs.

Here we investigated the genetic diversity of the growing number of phages belonging to the *Agtrevirus* genus and further characterized the host binding capabilities of agtrevirus phage AV101. Whole-genome comparison of 14 phages in the *Agtrevirus* genus showed that the *tsp* gene cluster represented the most diverse region. *In-silico* analysis of the TSPs showed that the receptor binding domains of phage AV101 were not observed in any *Agtrevirus* phages, but similar receptor binding domains can be found in other phages and prophages. We further determined the host recognition of the four TSPs which correlates with the O-antigens of the bacterial hosts. Our work expands the understanding of TSP host recognition in the *Agtrevirus* genus and give new insight into exchange of receptor binding domain between phages in different families and genera.

## Results

### Comparative genomics of *Agtrevirus* phages

*Agtrevirus* are known to infect diverse species within the *Enterobacteriaceae* family like *Shigella*, *Salmonella, Enterobacter* and *E. coli* including ESBL (Akter *et al*. 2019; Thanh *et al*. 2020; Kwon *et al*. 2021; Imklin *et al*. 2022; Vitt *et al*. 2023). To investigate the genetic differences that may account for such diverse host ranges, we extracted all *Agtrevirus* genomes as well as unclassified *Aglimvirinae* genomes, not to miss any potential *Agtrevirus* phages (Table S1). We did not include the unclassified *Aglimvirinae Dickeya* phages: phiDP10.3 and phiDP23.1 as they have previously been suggested to belong to the *Limestonevirus* genus in the *Aglimvirinae* subfamily (Czajkowski *et al*. 2015). We confirmed that all phages in the genus as well as the unclassified phages belonged to *Agtrevirus* by aligning all genomes and demonstrate an overall average nucleotide identity (ANI) between 86-97% (Figure S1). As phages PH4 and PC3 are 99.99 % identical, the same phage may have been isolated twice. Phylogenetic analysis using whole genome sequences demonstrated grouping of phages infecting *Salmonella* and *Shigella*, whereas *E. coli* phages formed a separate group with *Enterobacter* phage fGh-Ecl02. In contrast, *Enterobacter* phage EspM4VN and *Salmonella* phage SNUABM-02 did not group with any other phages (Figure 1A).

**Figure 1:**
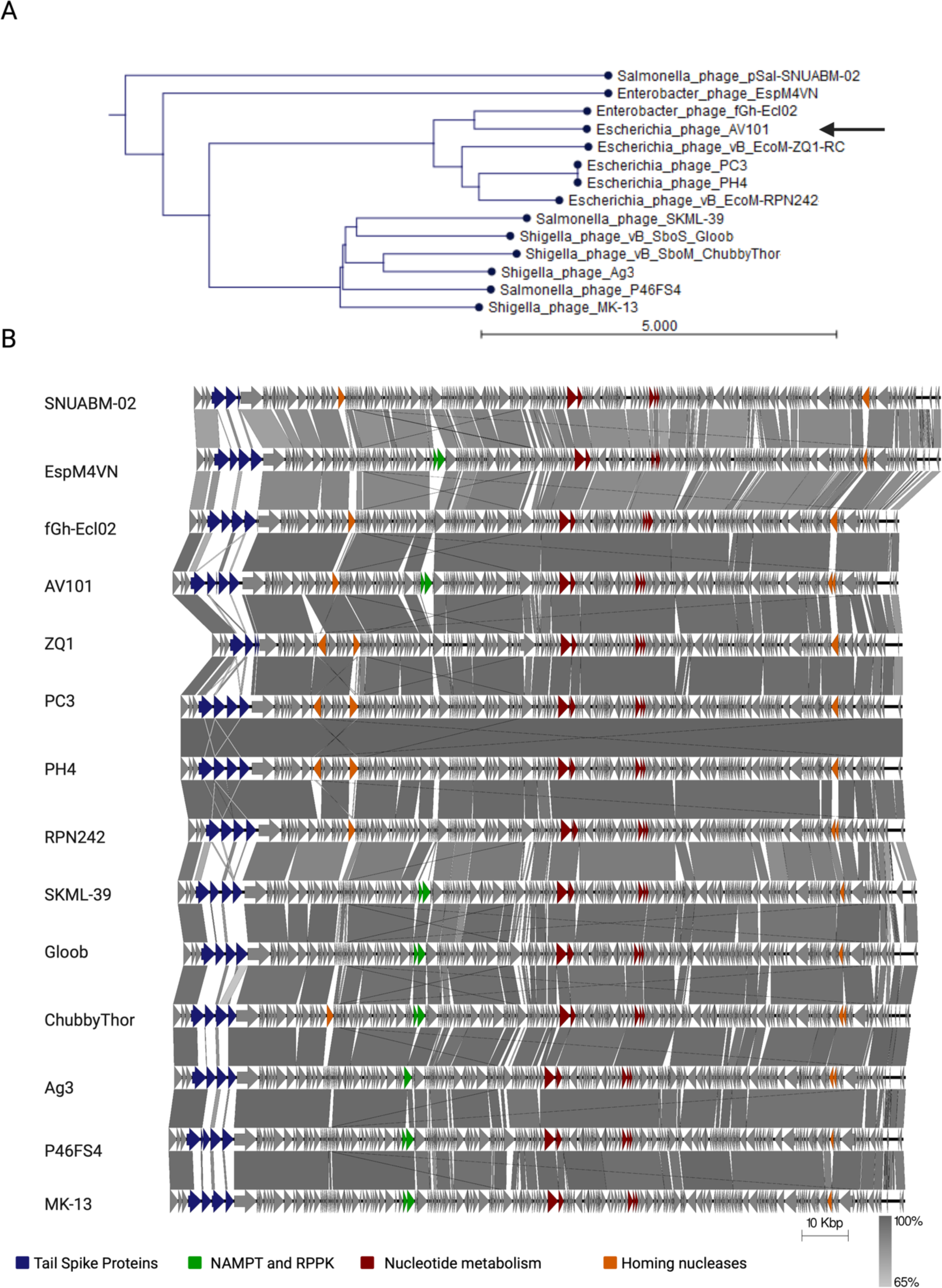
*Agtrevirus* phages are genetic similar with a few expectations. A) Average nucleotide identity phylogenetic analysis of phage AV101 and the other 13 extracted *Agtrevirus* phages using CLC workbench 22. The analysis showed that AV101 closes relative is Enterobacter phage fGh-Ecl02. B) Based on the phylogenetic analysis the *Agtrevirus* genomes were aligned using EasyFig. Genetic differences are highlighted with colours; Genes encoding nicotinamide phosphoribosyl transferase (NAMPT) and ribose-phosphate pyrophosphokinase (RPPK) (green), other nucleotide metabolism genes (red), homing nucleases (orange) and the *tsp* gene cluster (blue). Phage genomes and the variable genes are listed in supplementary table S2.

To further investigate the genetic differences of the *Agtrevirus* phages, we aligned all genomes using EasyFig (Sullivan, Petty and Beatson 2011). Overall, the phage genomes showed similar genetic organisation, and all encode hydroxymethyluracil (HMdU) transferase genes used for synthesis of hydroxylmethyl uracil replacing thymine as previously described (Figure 1B and Table S2) (Adriaenssens *et al*. 2012b, 2012a; Hsu *et al*. 2013). Interestingly, other nucleotide metabolism genes like nicotinamide phosphoribosyl transferase (NAMPT) and ribose-phosphate pyrophosphokinase (RPPK) were detected in eight of the 14 phage genomes analysed (Figure 1B and Table S2). These genes were associated with the group of phages classified as *Salmonella* and *Shigella* phages, but also in *E. coli* phage AV101 and *Enterobacter* phage EspM4VN. Beside genes involved nucleotide metabolism, we observed diversity within genes encoding homing nucleases, yet with no corelation to the phylogenetic groups (Figure 1B and Table S2). Several other genes varied between phages, but all were annotated as hypothetical proteins. Finally, the gene cluster encoding tail spike proteins expected to be responsible for host recognition exhibited the highest diversity (Figure 1B). Our overall genomic analysis showed that the *Agtrevirus* phages grouped into two phylogeny groups with two outliners and that the phages are highly conserved, except for the *tsp* gene cluster.

### *Agtrevirus* phages encode highly diverse tail spike proteins

To further analyse the diversity of TSPs of *Agtrevirus* phages, we compared the *tsp* gene cluster of the 14 phages (Figure 2A). While most *Agtrevirus* phages, like AV101, encode four *tsp* genes, phage ZQ1 and pSal-SNUABM-2 only encode two *tsp* genes. In phage AV101, we further confirmed the presence of multiple TSPs by performing transmission electron microscopy showing the expected star-like complex protruding the baseplate (Figure 2B).

**Figure 2:**
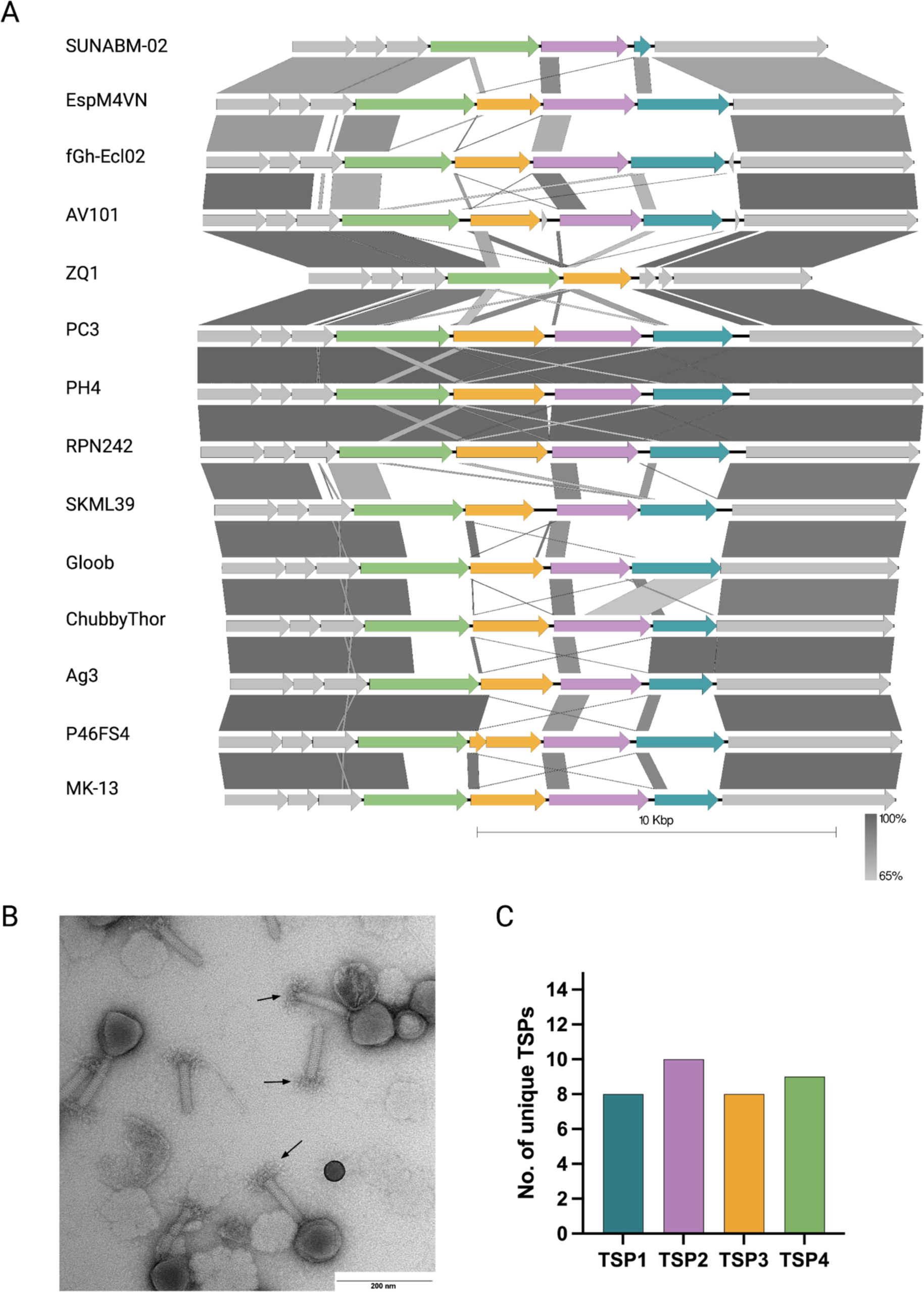
The majority of *tsp* genes in *Agtrevirus* phages encode unique receptor binding domains. A) Alignment of the *tsp* gene cluster located between the Virulence associated gene (*VriC)* and baseplate wedge genes (grey) of all available *Agtrevirus* phages showed high diversity of the *tsp* genes among the phages. Colours; Turkish: TSP1, purple: TSP2, yellow: TSP3, green: TSP4 and grey: hypothetical proteins, baseplate wedge and *VriC*. B) TEM photo of AV101 showing virion morphology similar to other *Agtrevirus* phages. The arrows points to the distinctive star-like TSP complex. C) All *tsp1, tsp2, tsp3* and *tsp4* genes were individually aligned and the number of dissimilar genes of *Agtrevirus tsp* genes were visualized plotted.

Analysing the *tsp* sequences in more detail demonstrated that most *tsp* genes were similar in the N-termini, which are known to be conserved within phage genera due to their importance for the TSP complex formation and hinging to the baseplate (Plattner *et al*. 2019; Chao *et al*. 2022) (Figure 2A). Furthermore, we observed a small region of nucleotide identity in the N-termini immediately upstream the β-helix responsible for receptor binding in *tsp1*, *tsp3* and *tsp4* genes of some phages (Figure 2A). This small region of nucleotide identity coincides with the tandem repeat domain previously suggested as a location for recombination between receptor binding domains of *tsp* genes of *Kuttervirus* phages (Sørensen *et al*. 2021).

Even though comparative genomics revealed two groups of *Agtrevirus* (Figure 1A), no apparent correlation between *tsp* genes and phylogenetic groups were observed. For instance, the group of *E. coli* phages encode diverse *tsp* genes expect from phage PH4 and PC3 (99.9% identical) and phage RPN242 that encode identical *tsp* clusters (Figure 1A, 2A and S1). In the group of *Shigella* and *Salmonella* phages, *Shigella* phage Ag3 encode *tsp4* with similarity to *Salmonella* phage P46FS4 and *tsp1* with similarity to *Shigella* phage ChubbyThor (Figure 2B). Interestingly, the region coding for the receptor binding domain of ChubbyThor *tsp2* was similar to the receptor binding domain of *Shigella* phage Gloob *tsp1* gene (Figure 2A). Thus, within this group, some of the *tsp* genes are similar but overall, most of the receptor binding domains of the *tsps* are diverse. Furthermore, aligning all *tsp1*, *tsp2*, *tsp3*, *tsp4* genes individually confirmed that *Agtrevirus* phages mainly express receptor binding domains unique to each phage, thus suggesting a total of 35 different receptor recognitions (Figure 2C). In summary, our analysis showed that most *Agtrevirus* phages encode *tsp* genes that only show N-terminus sequence similarity.

### The tail spike proteins of AV101 recognize specific O-antigens of ESBL *E. coli*

### hosts

The unique receptor binding domains of *Agtrevirus* TSPs suggest that they recognize different bacterial receptors. To further investigate host recognition of *Agtrevirus*, we used AV101 as an example. AV101 was previously shown to infect 9 out of 198 ESBL *E. coli* strains tested (Vitt *et al*. 2023).To further investigate the host range, we spotted a serial dilution of phage AV101 on the *E. coli* ECOR strain collection (n=72) as well as representative strains of *Salmonella enterica* subspecies Derby, Typhimurium, Enteritidis, Seftenberg, Anatum, Odersepoort and Minnesota. None of the tested strains could be infected by phage AV101, as no single plaques could be observed by performing a standard plaque assay (data now shown). The host range of AV101 was thus limited to the previous identified nine ESBL *E. coli* hosts (Figure 3A and 3B).

**Figure 3:**
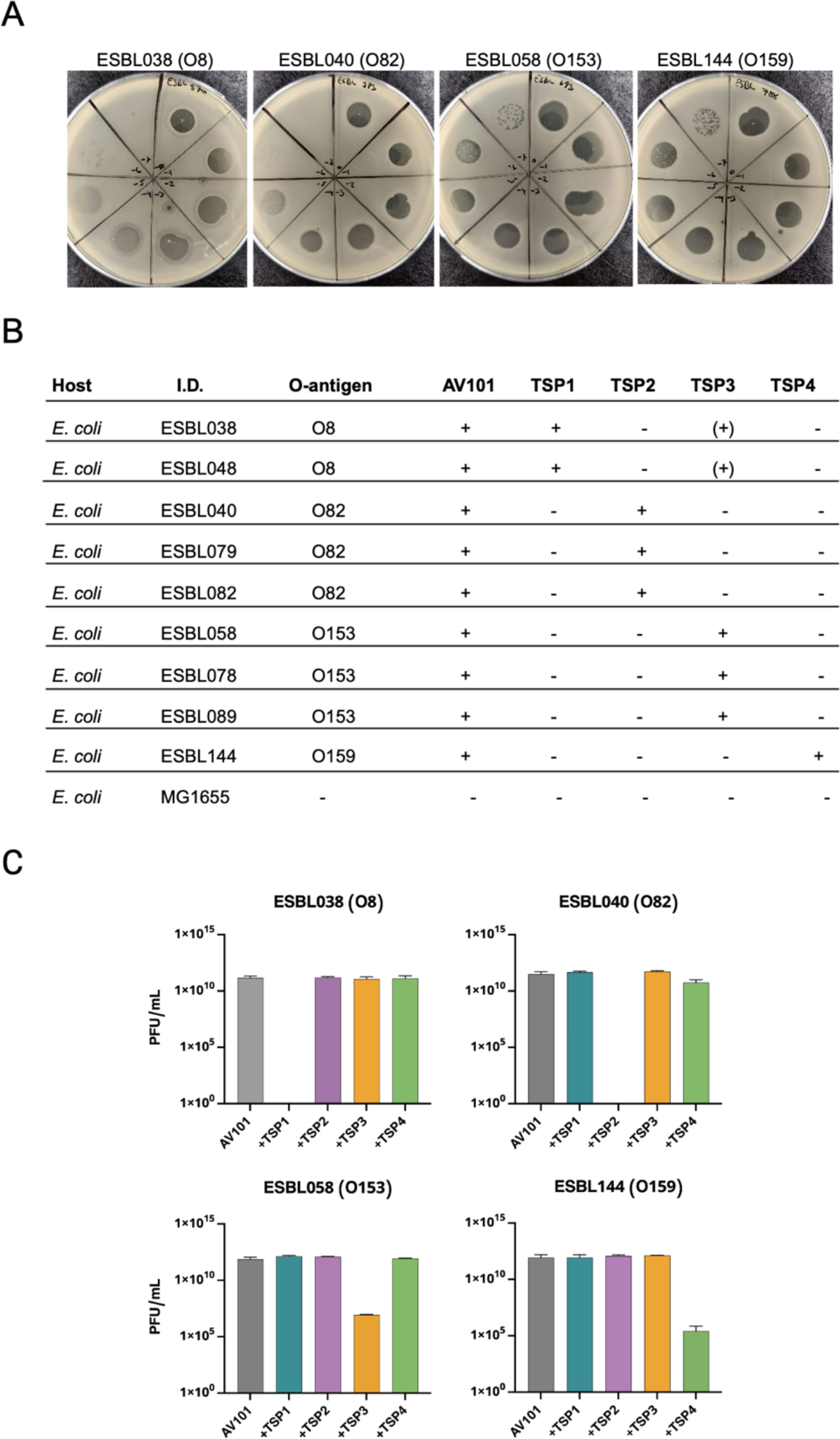
The four TSPs of AV101 recognize different O-antigens of ESBL *E. coli*. A) Phage AV101 infection on different bacterial hosts showing diverse plaque morphologies. B) Successful phage infection or detection of a clear translucent zone on bacterial lawns are indicated with plus sign. TSP3 makes small translucent zones on two *E. coli* strains expressing the O8 O-antigen (+). Big clear translucent zone is indicated with a + sign. Small translucent zone is indicated with (+) sign. C) Inhibition of AV101 infectivity on ESBL *E. coli* host after individual TSPs was preincubated with the ESBL hosts. TSP1 and TPS2 were able to completely block the infection of AV101 on their respective hosts, whereas only a partial inhibition with TSP3 and TSP4 could be observed.

To identify the receptors of the individual TSPs encoded by Agtrevirus phage AV101, we cloned, expressed, and purified the four TSPs and spotted each of them on the nine known ESBL *E. coli* hosts. After incubation we noted if the TSPs were able to degrade the O-antigen by forming translucent zones on the bacterial lawns (Figure 3B and Figure S2). By this analysis, we observed a correlation between the TSP type and the O-antigen encoded by the individual bacterial hosts as TSP1, TSP2, TSP3, and TSP4 formed translucent zones only on O8, O82, O153 and O159 ESBL *E. coli* strains, respectively (Figure 3B). Yet, TSP1 were not able to make translucent zones on the *E. coli* strains ECOR7 and ECOR72 expressing O8 O-antigen (data not shown). Surprisingly, TSP3 also formed a translucent zone on O8 hosts although smaller compared to TSP1 (Figure S2), suggesting that TSP3 may be able to degrade both O153 and O8 antigens.

To show that binding of the individual TSPs to specific *E. coli* O-antigens are important for AV101 infection, we carried out an inhibition assay. For this experiment, we chose ESBL038 (O8), ESBL040 (O82), ESBL058 (O153) and ESBL144 (O159) to represent hosts recognised by TSP1, TSP2, TSP3 and TSP4, respectively. The strains were grown to exponential phase and mixed with the individually TSPs, allowing the TSPs to bind to and degrade their receptor before plating to form a lawn. Afterwards, AV101 were spotted on the lawn to evaluate the ability of the phage to access the receptor and subsequently form plaques. TSP1 and TSP2 completely abolished the infection of AV101 on ESBL038 and ESBL040, respectively (Figure 3C). In contrast, TSP3 and TPS4 had an inhibitory effect on infection of ESBL058 and ESBL144, respectively, leading to approximately 5-log reduction of the phage titre (Figure 3C). The differences in the ability of the TSPs to inhibit infection may suggest different kinetics of the enzymatic activity of the receptor binding domains. Finally, while we observed small translucent zones on ESBL038 (O8 O-antigen) when spotting TSP3, the protein did not inhibit infection of phage AV101 of ESBL038. This suggest that TSP3 does not degrade the O8 O-antigen but may degrade another surface polysaccharides of this strain, thus forming a translucent zone unrelated to O-antigen degradation. Overall, our results demonstrate phage AV101 express four TSPs that are unique for the *Agtrevirus* genus and each of the TSPs recognize distinct O-antigens required for infection of ESBL *E. coli*.

### The receptor binding module of phage AV101 tail spike proteins show similarity to prophages and virulent phages

While we only observed little *tsp* sequence similarity between AV101 and other *Agtrevirus* phages, it is known that the receptor binding domain of TSPs may be subjected to horizontal gene transfer between phages from distant related families (Pires *et al*. 2016; Latka *et al*. 2019; Sørensen *et al*. 2021). Thus, to further investigate if the receptor binding domain could be found in other phages than *Agtrevirus*, we extracted the amino acid sequence and conducted a BLASTp analysis of the four TSPs of phage AV101. TSP1, TSP2 and TSP3 did not show overall similarity to any virulent phage genomes except for the conserved N-termini of TPSs of *Ackermannviridae* phages (data not shown). Instead, similarities in the C-terminal were found to *E. coli* genome sequences, suggesting that these TSPs originate from diverse prophages (Figure 4A and Table S3). To identify and further classify the corresponding prophages, we extracted and analysed these *E. coli* genomes using PHAge Search Tool Enhanced Release (PHASTER)(Arndt *et al*. 2016). Indeed, the analysis identified prophages in all the *E. coli* genomes that were either intact or questionable (Table S4), suggesting that these prophages express TSP with similar receptor binding domains. Furthermore, we investigated if the TSPs shared receptor binding domains in prophages encoded by our own ESBL *E. coli* strains collection (n=198). However, none of the four TSPs shared similarity to prophages in our collection (data not shown). In contrast, TSP4 showed similarity to virulent phages infecting *E. coli* as well as *Salmonella* including kuttervirus LPST94, kayfunavirus ST31 and phapecoctavirus Ro121c4YLVW (Figure 4B). The phages ST31 and Ro121c4YLVW infects *E. coli* and are only distantly related to *Agtrevirus*, whereas LPST94 belongs to *Ackermannviridae* and infects *Salmonella enterica* subspecies (Liu *et al*. 2018; Yan *et al*. 2020; Khalifeh *et al*. 2021). Still, the receptors recognized by these TSPs have not been identified (Yan *et al*. 2020). To get a better understanding of the similarity in relation to the structural domains, we used Alphafold2-multimer to predict the structure of the four TSP of AV101 (Figure 4B). All the TSPs shared a modular fold, with an anchor domain in the N-terminus, and a receptor binding domain close to the C-terminal. Beside the β-helix, the N-terminal head-binding domains of the TSPs had a low prediction score (Figure S3 and S4). When we compared the alignment of TSPs and the structures, we observed that the amino acid similarity coincides with the β-helix carrying the receptor binding domain (Figure 4A and B). Thus, we expect that the prophages and virulent phages bind to the same O-antigen receptors as the TSPs of phage AV101.

**Figure 4:**
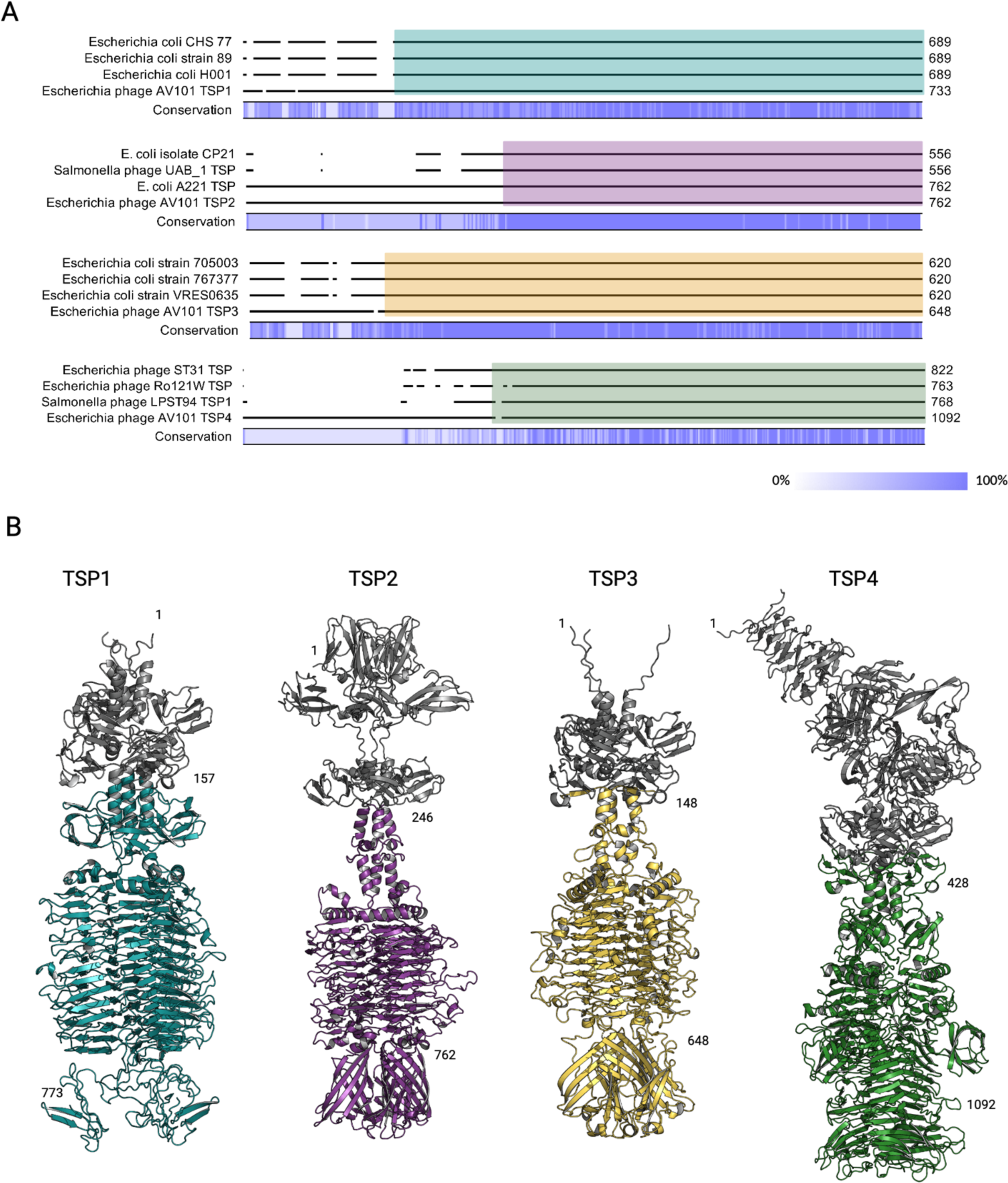
The receptor binding domain of AV101 TSPs show similarity to prophages and distant related lytic phages. A) BLASTp analysis of the four TSPs showed that TSP1, TSP2 and TSP3 share similarity towards proteins found in E. coli strains. Furthermore, TSP2 and TSP4 had similarity towards distant related lytic phages. A detailed overview of the sequence similarity and the accession numbers of the BLASTp hits can be found in supplementary table S3. The coloured boxes represent the region of similarity visualized in the structures. B) AlphaFold 2 was used to prediction and visualize the four TSPs of AV101. The four TSPs folds like other known TSPs with the conserved β-helix serving as the receptor binding domain. Moreover, the analysis showed that the similarity to other TSPs were found in the receptor binding domains, thus are highlighted in the same colors as the alignment.

## Discussion

With the rise of genome-based phage taxonomy, the investigation of biological functions of a single phage can provide a general understanding of other phages within the same family or genus (Turner, Kropinski and Adriaenssens 2021). *Agtrevirus* phages infect different *Enterobacteriaceae* species, but not much is known about the genetic differences between the phages allowing them to infect such diverse hosts (Akter *et al*. 2019; Thanh *et al*. 2020; Kwon *et al*. 2021; Imklin *et al*. 2022). Receptor-binding proteins are responsible for the initial binding to the bacterial hosts and the characterization of such proteins can provide important biological information, crucial for understanding phage host ranges. Here we investigated phages belonging to the *Agtrevirus* genus and observed that while all phages show high nucleotide identity and a similar genome structure, the genes encoding tail spike proteins (TSPs) were highly diverse. Furthermore, as an example, we investigated the specific host recognition of the four TSPs encoded by *Agtrevirus* phage AV101 infecting ESBL *E. coli*.

To investigate similarities and difference among *Agtrevirus* phages, we carried out a comparative *in-silico* analysis of all available *Agtrevirus* genomes. Besides observing diversity in the *tsp* gene region, we also observed that eight of the phages encoded nicotinamide phosphoribosyl transferase and ribose-phosphate pyrophosphokinase. These genes have also been observed in distant related phages including *Babavirus* Baba19a, *Schizotequatrovirus* KVP40 and polybotosvirus Atu_ph7 phages (Lee, Li and Miller 2017; Attai *et al*. 2018; Nilsson *et al*. 2019). Ribose-phosphate pyrophosphokinase convert ribose 5-phosphate and ATP to phosphoribosyl pyrophosphate (PRPP) and AMP. Phosphoribosyl pyrophosphate is a precursor of purine and pyrimindine that is used by ribonucleotide reductases, also encoded by all *Agtrevirus* phages, for nucleotide production during the phage replication (Nilsson *et al*. 2019). Given that not all *Agtrevirus* phages encode these genes, they may not be required for phage genome replication but provide other advantages, yet to be discovered, during certain conditions or when infecting specific hosts.

Previously, we have characterized TSPs encoded by phages of the *Ackermannviridae* family and observed that receptor binding domains were conserved between phages especially within the *Kuttervirus* genus (Sørensen *et al*. 2021). In contrast, we showed that the receptor binding domain are not as well conserved within the *Agtrevirus* genus, as most phages encode unique domains, indicating specific host recognition. Instead, our protein alignment and AlphaFold2 predictions showed that the receptor binding domain of phage AV101 TSPs are similar to distant related virulent phages as well as prophages. Likewise, the receptor binding domain of TSP1 of kuttervirus CBA120 show similarity to TSP encoded by a prophage found in a *Salmonella* Minnesota strain (Plattner *et al*. 2019) and similar receptor binding domains were identified in kuttervirus Det7 and temperate lederbergvirus P22 (Walter *et al*. 2008). Thus, more receptor binding domains of TSPs found in the *Ackermannviridae* family may originate from prophages. Furthermore, the receptor binding domain of TSP4 of AV101 was similar to TSP1 encoded by phage *Kuttervirus* LPST94, demonstrating exchange of receptor binding domains between phages of different genera within the *Ackermannviridae* family. So far, host range analysis demonstrated that phage LPST94 infects several *Salmonella enterica* subspecies expressing different O-antigens (Yan *et al*. 2020), yet phage AV101 could not infect any of these *Salmonella enterica* subspecies. In addition, phage LPST94 could infect six *E. coli* strains, but information about their O-antigens was not included in the study (Yan *et al*. 2020). Still, our study suggests that TSP1 of phage LPST94 may recognize O8 O-antigen, but it remains to be experimental verified. Overall, combining bioinformatic, structural predictions and biological data of phage AV101 TSPs gives an insight into the origin and acquisition of the receptor binding abilities of *Ackermannviridae* phages.

ESBL *E. coli* are of high concern as they encode extended spectrum β-lactamases conferring resistance to antibiotics such as penicillin and cephalosporins commonly used in treatment human infections (Paterson and Bonomo 2005; Benz *et al*. 2021). To propose alternative solutions targeting ESBL *E. coli*, we recently established a collection of phages, including phage AV101, infecting ESBL *E. coli* (Vitt *et al*. 2023). Using this collection, we composed phage cocktails that prevented growth of ESBL *E. coli*, thus suggesting that phages may indeed be promising alternative antimicrobials (Vitt *et al*. 2023). Phage AV101 was not included in these cocktails, but may be used to target ESBL *E. coli* expressing O8, O82, O153 and O159 O-antigens due to the specificities of the four TSP. It should be noted though, that AV101 did not infect two other *E. coli* strains tested (ECOR7 and ECOR72) carrying the O8 O-antigen, suggesting that internal defence mechanism or modification of the O-antigen, like glycosylation or acetylation may influence phage infection (Knirel *et al*. 2015; Egido *et al*. 2022). In general, the four O-antigens are expressed in diverse pathogenic and commensal *E. coli* strains (Tamaki *et al*. 2005a; Marin *et al*. 2022). For example, O159 O-antigen are often expressed by enterotoxigenic *E. coli* (ETEC) causing diarrhoea in humans (Linnerborg, Weintraub and Widmalm 1999; Tamaki *et al*. 2005b), suggesting that AV101 may be used in phage therapy or biocontrol of such strains. Thus, identification of receptor recognition of AV101 may allow design of phage applications dedicated to pathogenic *E. coli* like O159 ETEC or ESBL *E. coli*.

Purified TSPs have also demonstrated promising potential as therapeutics or diagnostic tools (Wang *et al*. 2023). For instance, oral administration of purified TSP of *Lederbergvirus* phage P22 dramatically decreased *Salmonella* colonization in the chicken gut and reduced penetration into internal organs (Waseh *et al*. 2010). TSPs from bacteriophages 9NA and P22 have been used as tools to detect *S. typhimurium* (Schmidt *et al*. 2016) and TSP3 of kuttervirus Det7 was used as a biosensor using surface plasmon resonance to detect *S. typhimurium* (Hyeon, Lim and Shin 2020). Thus, instead of using phage AV101 as an antibacterial agent, AV101 TSPs targeting the O-antigens on ESBL *E. coli* or O159 of ETEC strains could be of interest. Furthermore, other phage-based solutions for combating pathogenic bacteria based on exploring the knowledge of binding abilities TSPs proteins may be developed. For example, pyocins are phage tail-like particles encoded by *Pseudomonas aeruginosa* showing bactericidal activity against other *Pseudomonas* strains (Ge *et al*. 2020). Such pyocins have been engineered to successful kill other bacteria by exchanging the receptor binding protein of the native pyocin with phage receptor binding proteins (Williams *et al*. 2008; Scholl *et al*. 2009). Similarly, TSPs targeting specific O-antigens, like those encoded by AV101, may be used to re-target pyocins to other hosts. TSPs may also be utilized for developing Innolysins, which are novel antibacterials fusing a receptor binding protein or domain to an endolysin, allowing the engineered enzyme to target gram negative bacteria (Zampara *et al*. 2020). So far Innolysins have been created and shown to kill specifically *Campylobacter jejuni* or commensal *E. coli* (Zampara *et al*. 2020, 2021), and the use of TSP4 could allow development of novel Innolysins targeting *E. coli* O159 strains including ETEC. Thus, investigation of the receptor recognition of TSPs is not only crucial for understanding the host range of phages but can be further exploited for the development of therapeutics against pathogenic bacteria.

## Material and methods

### Bacterial strains and phages

All phage genomes used for genomic analysis are presented in supplementary Table S1. A list of the bacterial strains used in the study is presented in supplementary Table S4.

### Bioinformatic analysis

All genomes of phages from the *Agtrevirus* genus and unclassified *Aglimvirinae* subfamily were extracted from NCBI. Phylogenetic analysis of all *Agtevirus* phages was performed using whole genomes sequence in CLC genomics version 22 (Qiagen) with default settings (date 04/01-23). To align and visualize all the genomes as well as the *tsp* gene cluster Easyfig version 2.2.5 or CLC genomics were used (Sullivan, Petty and Beatson 2011). 0.4 minimum identity was chosen as the BLAST setting. To identify homolog TSPs, BLASTp was used with standard settings. Afterwards, the genomes identified was used to analysis of prophage in the *E. coli* genomes and was carried out with PHASTER with standard settings (Arndt *et al*. 2016).

### Phage propagation

Agtrevirus AV101 was propagated on *E. coli* strain ESBL058 as described earlier (Sørensen *et al*. 2021). A single colony of *E. coli* ESBL058 was inoculated into LB media (Lysogeny Broth, Merck, Darmstadt, Germany) and incubated until exponential phase at 37 °C at 180 rpm. A previous phage stock of AV101 (1.6×10^10^ PFU mL^−1^) was 10-fold diluted followed mixing 100 μL of the dilutions with 100 μL of the ESBL058. On a LA plate (LB with 1,2% agar), 4 mL of molten top agar (LBov; LB broth with 0,6% Agar bacteriological no.1, Oxoid) was applied. 5 mL of SM buffer (0.1 M NaCl, 8 mM MgSO47H2O, 50 mM Tris-HCl, pH 7.5) was added to the plates after an overnight incubation at 37 °C, and the plates were then incubated at 4 °C at 50 rpm. The overnight samples were collected, centrifuged for 15 minutes at 11000 rpm, then passed through a 0,2 M filter. By using a phage plaque assay (described below), the new phage stock was used to estimate the phage titre.

### Phage DNA isolation

AV101 phage DNA was extracted using phenol-chloroform as described earlier (Gencay *et al*. 2019). Briefly, 1 mL phage lysate was filtered three times with 0.22 µM syringe filters. RNase (10 µg mL^−1^) and DNase (20 µg mL^−1^) were added to the filtered lysate and incubated for one hour at 37°C. The degradation was stopped by adding sterile EDTA (pH 8) at a final concentration of 20 mM. Afterwards Proteinase K was added (50 µg mL^−1^) and incubate for two hours at 56°C followed by cooling the sample to room temperature. To isolate the phage DNA, we used Genomic DNA clean and concentrator™ (Zymo research) following the manufactures instructions. The DNA concentration was measured using Qubit 2.0 Fluorometer (Invitrogen).

### Transmission electron microscopy

To visualize phage AV101 with transmission electron microscopy we used a previously described method (Ackermann 2009). Briefly, bacteriophages from high-titer stock were sedimented at 12000g for 60 min at 4 C° and washed three times with Ammonium Acetate (0.1 M, pH 7). Final sediment was used for imaging. 200 mesh copper coated carbon grids (Ted Pella, Inc.) were made hydrophilic by glow discharging the grids using a Leica Coater ACE 200 for 30 sec at 10 mA. 6 µL of phages at a PFU mL^−1^ of 10^12^ were pipetted on the grids and incubated for 30 seconds. All liquid was removed with a Whatman filter paper. The phages were stained by incubating the grid with 6 µL of 2% uranyl acetate for 30 seconds. In a washing step, 6 µL of ddH2O was pipetted on the grid, incubated for 30 seconds, and removed with a Whatman paper. The phages where imaged using a CM100 microscope with a Bio TWIN objective lens and a LaB6 emitter. Pictures were taken with an Olympus Veleta camera and analyzed using ImageJ to determine phage particle measurements.

### Phage host range analysis

The newly prepared phage stock (3.4×10^12^ PFU mL^−1^) was used to determine the host range of AV101 (Gencay *et al*. 2019). Bacterial strains were inoculated into 5 mL LB medium (Lysogeny Broth, Merck, Darmstadt, Germany) and grown for 5 hours. Afterwards, 100 µL of the strain were suspended into 4 mL of top-agar and poured onto LB plates left to solidify (approximately 30 minutes). 10-fold serial dilutions of the phage stock were prepared, and three times of 10 µL of each dilution were spotted on the bacterial lawn. Subsequently the spots were dried the plates were incubated overnight at 37°C. The next day, dilutions with single plaques were counted to calculate the PFU mL^−1^.

### Tail spike protein cloning

Purified AV101 genomic DNA was used as a template for *tsp* cloning using In vivo assembly cloning (García-Nafría, Watson and Greger 2016; Sørensen *et al*. 2021). The tsp genes (*tsp1-* (2202 bp.), *tsp2-* (2289 bp.), *tsp3-* (1947 bp.) and *tsp4-* (3279 bp.)) were individually amplified with Phusion™ High-Fidelity DNA polymerase (Thermo Scientific™) following the manufacture instructions with primers carrying homologous overhangs to the expression vector pET-28a (+). Furthermore, the expression vector with primers to linearize the vector were also added to each PCR reaction. The primers amplifying the *tsp* genes would allow for homologous recombination between the HinCII and Eco52KI restriction sites in the multiple cloning site of the vector. Primers (Table 1) were ordered from TAG Copenhagen A/S. Furthermore, the cloning site in the pET-28a (+) also express a his-tag upstream so the expressed TSPs could be purified with affinity chromatography. In order to degrade the parental pET28a-(+) plasmid, 1 µL of FastDigest DpnI enzyme (Thermo ScientificTM) was added to the PCR reactions after the amplification of the *tsp* genes and the expression vector. Each of the PCR samples were transformed into competent E. coli StellarTM cells (Takara Bio). The next day, pET_AV101_TSP1-4 extracted with GeneJET Plasmid Miniprep Kit (Thermo Scientific™). Vectors carrying the correct *tsp* insert (pET_AV101_TSP1-4) were confirmed by Sanger sequencing (Eurofins Genomics).

**Table 1:**
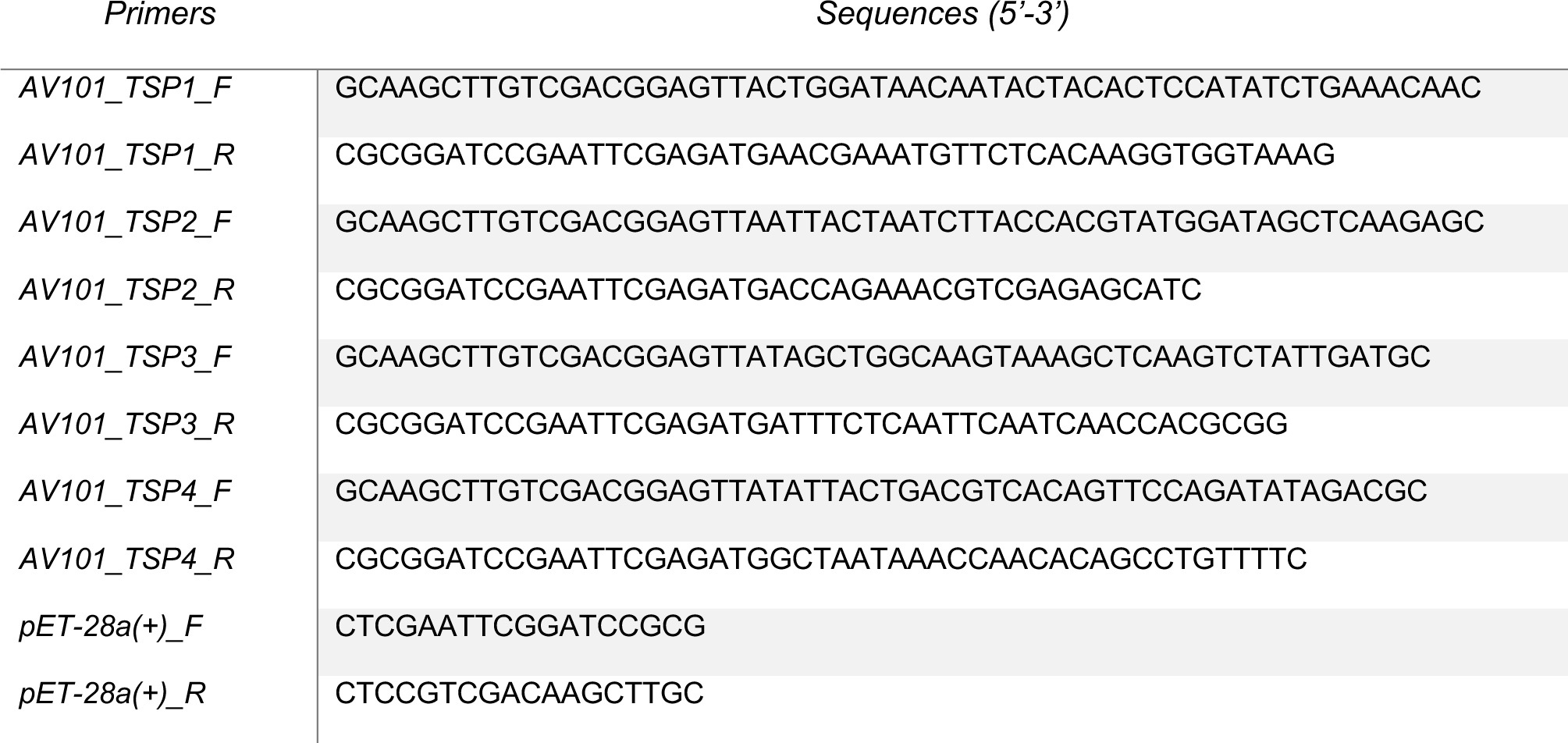
overview of primers for cloning of the four *tsp* genes.

### Tail spike protein purification

pET_AV101_TSP1-4 were transformed into electrocompetent *E. coli* BL21 cells for the expression of the TSPs (Sørensen *et al*. 2021). A single colony of each of transformants carry one of the four pET_AV101_TSP1-4 was inoculated into LB medium with 50 µg mL^−1^ kanamycin and incubated overnight at 37°C and 170 rpm. Next day, 10 mL of the starter culture was added to 1L of LB medium with 50 µg mL^−1^ kanamycin at 37°C and incubated until an OD_600_ value of 0.6. Then, a final concentration of 0.5 mM of isopropyl-b-D-thiogalactopyranoside (IPTG) was added to induce protein expression. The culture was incubated for an entire night at a reduced temperature of 16°C. The culture was centrifuged the following morning for 10 minutes at 13000 g, and the pellet was then resuspended in 9 mL of lysis buffer (0.5 M NaCl, 20 mM Na2HPO4, 50 mM Imidazole, pH 7.4). Sonication was used to disrupt the cells, with a program of 9 cycles lasting 30 seconds each at 80% power. Centrifugation at 9500 g for 30 min. at 4°C separated cell debris. Using 0.22 mm filters, the supernatant containing the expressed proteins was filtered. HisGraviTrapTM (GE Healthcare) was then used to purify the proteins using an elution buffer (0.5 M NaCl, 20 mM Na_2_HPO_4_, 0.5 M Imidazole, pH 7.4). To transfer the TSPs into a new buffer (20 mM HEPES (pH 7.4)), Amicon Ultra-15 Centrifugal Filter Units with a 50 kDa cutoff (Merck Milipore) were utilized. Using a Qubit 2.0 Fluorometer and the QubitTM Protein Assay Kit, protein concentration was determined (Invitrogen).

### Tail spike protein spot assay and inhibition assay

The TSP spot and inhibition assay was performed as previously described (Sørensen *et al*. 2021). Shortly, bacterial strains were grown to an OD_600_ value of 0.6. 100 µL of the bacterial culture was then added to 4 mL top-agar and poured onto an LB agar plate. After the bacterial lawns were solidified, 1.5 µg of the four TSPs were spotted onto the bacterial lawn and left to dry for 30 minutes. Phage AV101 and the protein buffer (20 mM HEPES (pH 7.4)) were used as a positive and negative control, respectively. The plates were incubated overnight at 37°C and the next day the presence of a translucent zone was evaluated. To further validate the spot assay, the inhibitory effect of the TSPs on the infectivity of the bacterial strains were evaluated. The four strains ESBL-038, -040, -058, -144 were used as they represented strains that TSP1-4 recognize, respectively. A single colony of the bacterial strains was and incubated in LB medium at 37°C at 170 rpm until OD_600_ reached 0.3. The cells were then cooled on ice before 100 µL of the cells were added to 5 mg of the TSPs in individually tubes. The cell-TSP suspension was then preincubated at 37 °C for 20 min. Afterwards, the suspension was added to 4 mL top agar and poured onto an LB agar plate and left to solidify. Three times 10 µL phage AV101 dilutions (10^−1^ to 10^−8^) was spotted on top of the plate and incubated overnight at 37°C. Next day, the inhibitory effect of the TSPs were evaluated by comparing the PFU mL^−1^ of the phage sensitive strains with the PFU mL^−1^ with strains incubated with the respective TSPs. The inhibition assay was carried out in triplicates and the results were visualized in Graphpad Prism9 with the mean standard deviations shown in the figure.

### Alphafold2 prediction

Models for the TPSs were predicted in a Nvidia Quadro RTX 8000 using Alphafold2-multimer (version 3.2.1) (Jumper *et al*. 2021; Evans *et al*. 2022). We utilized global search for the multiple sequence alignment, five recycling rounds, and the amber relaxation was skipped. The best model was ranked based on the iptm+ptm score and was the one selected for the analysis.

## Data availability

The genome sequence data of phage AV101 has been submitted to the National Center for Biotechnology Information (NCBI) (https://www.ncbi.nlm.nih.gov/nuccore/) under accession number OQ973471.

## Acknowledgements

This work was supported by the Danish Council for Independent Research (9041-00159B). We acknowledge the Core Facility for Integrated Microscopy, Faculty of Health and Medical Sciences, University of Copenhagen.

## CRediT authorship contribution statement

**Anders Nørgaard Sørensen:** Conceptualization, Methodology, Validation, Formal analysis, Investigation, Writing – original draft, Writing – review & editing. **Dorottya Kalmar:** Conceptualization, Methodology, Validation, Formal analysis, Investigation, Writing – review and editing. **Veronika Theresa Lutz:** Investigation, Formal analysis, Writing – review and editing. **Victor Klein-Sousa:** TSPs modelling, Writing – review & editing. **Nicholas M. I. Taylor:** Writing – review & editing. **Martine Camilla Holst Sørensen:** Conceptualization, Visualization, Writing – review & editing, Funding acquisition. **Lone Brøndsted:** Conceptualization, Project administration, Supervision, Visualization, Writing-review & editing, Project administration, Funding acquisition.

## Abbreviations

ESBL: Extended Spectrum β-lactamase
ETEC: Enterotoxigenic *E. coli*
HMdU: Hydroxymethyluracil
LB: Luria-Bertani
LPS: Lipopolysaccharide
NAMPT: Nicotinamide phosphoribosyl transferase
NCBI: National Center for Biotechnology Information
PFU: Plaque formation unit
RPPK: Ribose-phosphate pyrophosphokinase
TEM: Transmission electron microscopy
TSP: Tail spike protein
VriC: Virulence-associated protein

## References

Ackermann H-W. Basic Phage Electron Microscopy. In: Clokie Martha R.J. and Kropinski AM (ed.). Bacteriophages: Methods and Protocols, Volume 1: Isolation, Characterization, and Interactions. Totowa, NJ: Humana Press, 2009, 113–26.

Adriaenssens EM, Ackermann HW, Anany H et al. A suggested new bacteriophage genus: “Viunalikevirus.” Arch Virol 2012a;157:2035–46.

Adriaenssens EM, van Vaerenbergh J, Vandenheuvel D et al. T4-related bacteriophage LIMEstone isolates for the control of soft rot on potato caused by “Dickeya solani.” PLoS One 2012b;7, DOI: 10.1371/journal.pone.0033227.

Adriaenssens EM, Wittmann J, Kuhn JH et al. Taxonomy of prokaryotic viruses: 2017 update from the ICTV Bacterial and Archaeal Viruses Subcommittee. Arch Virol 2018;163:1125–9.

Akter M, Brown N, Clokie M et al. Prevalence of Shigella boydii in Bangladesh: Isolation and Characterization of a Rare Phage MK-13 That Can Robustly Identify Shigellosis Caused by Shigella boydii Type 1. Front Microbiol 2019;10, DOI: 10.3389/fmicb.2019.02461.

Anany H, Lingohr EJ, Villegas A et al. A Shigella boydii bacteriophage which resembles Salmonella phage ViI. Virol J 2011;8, DOI: 10.1186/1743-422X-8-242.

Andres D, Hanke C, Baxa U et al. Tailspike interactions with lipopolysaccharide effect DNA ejection from phage P22 particles in vitro. Journal of Biological Chemistry 2010;285:36768–75.

Arndt D, Grant JR, Marcu A et al. PHASTER: a better, faster version of the PHAST phage search tool. Nucleic Acids Res 2016;44:W16–21.

Attai H, Boon M, Phillips K et al. Larger than life: Isolation and genomic characterization of a jumbo phage that infects the bacterial plant pathogen, Agrobacterium tumefaciens. Front Microbiol 2018;9, DOI: 10.3389/fmicb.2018.01861.

Barbirz S, Müller JJ, Uetrecht C et al. Crystal structure of Escherichia coli phage HK620 tailspike: Podoviral tailspike endoglycosidase modules are evolutionarily related. Mol Microbiol 2008;69:303–16.

Benz F, Huisman JS, Bakkeren E et al. Plasmid- and strain-specific factors drive variation in ESBL-plasmid spread in vitro and in vivo. ISME J 2021;15:862–78.

Chao KL, Shang X, Greenfield J et al. Structure of Escherichia coli O157:H7 bacteriophage CBA120 tailspike protein 4 baseplate anchor and tailspike assembly domains (TSP4-N). Sci Rep 2022;12, DOI: 10.1038/s41598-022-06073-2.

Czajkowski R, Ozymko Z, de Jager V et al. Genomic, proteomic and morphological characterization of two novel broad host lytic bacteriophages PdblPD10.3 and PdblPD23.1 infecting pectinolytic Pectobacterium spp. and Dickeya spp. PLoS One 2015;10, DOI: 10.1371/journal.pone.0119812.

Egido JE, Costa AR, Aparicio-Maldonado C et al. Mechanisms and clinical importance of bacteriophage resistance. FEMS Microbiol Rev 2022;46, DOI: 10.1093/femsre/fuab048.

Evans R, O’Neill M, Pritzel A et al. Protein complex prediction with AlphaFold-Multimer. bioRxiv 2022:2021.10.04.463034.

García-Nafría J, Watson JF, Greger IH. IVA cloning: A single-tube universal cloning system exploiting bacterial In Vivo Assembly. Sci Rep 2016;6:1–12.

Ge P, Scholl D, Prokhorov NS et al. Action of a minimal contractile bactericidal nanomachine. Nature 2020;580:658–62.

Gencay YE, Gambino M, Prüssing TF et al. The genera of bacteriophages and their receptors are the major determinants of host range. Environ Microbiol 2019;21:2095–111.

Heyse S, Hanna LF, Woolston J et al. Bacteriophage cocktail for biocontrol of Salmonella in dried pet food. J Food Prot 2015;78:97–103.

Hsu CR, Lin TL, Pan YJ et al. Isolation of a Bacteriophage Specific for a New Capsular Type of Klebsiella pneumoniae and Characterization of Its Polysaccharide Depolymerase. PLoS One 2013;8, DOI: 10.1371/journal.pone.0070092.

Hyeon SH, Lim WK, Shin HJ. Novel surface plasmon resonance biosensor that uses full-length Det7 phage tail protein for rapid and selective detection of Salmonella enterica serovar Typhimurium. Biotechnol Appl Biochem 2020:1–8.

Imklin N, Sriprasong P, Thanantong N et al. Characterization and complete genome analysis of a novel Escherichia phage, vB_EcoM-RPN242. Arch Virol 2022;167:1675–9.

Jumper J, Evans R, Pritzel A et al. Highly accurate protein structure prediction with AlphaFold. Nature 2021;596:583–9.

Khalifeh A, Kraberger S, Dziewulska D et al. Complete Genome Sequence of a Phapecoctavirus Isolated from a Pigeon Cloacal Swab Sample. Microbiol Resour Announc 2021;10:e01471–20.

Knirel YA, Prokhorov NS, Shashkov AS et al. Variations in O-antigen biosynthesis and O-acetylation associated with altered phage sensitivity in Escherichia coli 4s. J Bacteriol 2015;197, DOI: 10.1128/JB.02398-14.

Kropinski AM, Anany H, Kuhn JH et al. 2017.001B.A.v1.Ackermannviridae. ICTV 2017.

Kunstmann S, Scheidt T, Buchwald S et al. Bacteriophage Sf6 tailspike protein for detection of shigella flexneri pathogens. Viruses 2018;10, DOI: 10.3390/v10080431.

Kutter EM, Skutt-Kakaria K, Blasdel B et al. Characterization of a ViI-like Phage Specific to Escherichia Coli O157:H7., 2011.

Kwon J, Kim SG, Kim HJ et al. Bacteriophage as an alternative to prevent reptile-associated Salmonella transmission. Zoonoses Public Health 2021;68:131–43.

Latka A, Leiman PG, Drulis-Kawa Z et al. Modeling the Architecture of Depolymerase-Containing Receptor Binding Proteins in Klebsiella Phages. Front Microbiol 2019;10, DOI: 10.3389/fmicb.2019.02649.

Lee IM, Tu IF, Yang FL et al. Structural basis for fragmenting the exopolysaccharide of Acinetobacter baumannii by bacteriophage ▪aB6 tailspike protein. Sci Rep 2017;7:1–13.

Lee JY, Li Z, Miller ES. Vibrio phage KVP40 encodes a functional NAD+ salvage pathway. J Bacteriol 2017;199, DOI: 10.1128/JB.00855-16.

Lefkowitz EJ, Dempsey DM, Hendrickson RC et al. Virus taxonomy: The database of the International Committee on Taxonomy of Viruses (ICTV). Nucleic Acids Res 2018;46:D708–17.

Linnerborg M, Weintraub A, Widmalm G. Structural studies utilizing 13C-enrichment of the O-antigen polysaccharide from the enterotoxigenic Escherichia coli 0159 cross-reacting with Shigella dysenteriae type 4. Eur J Biochem 1999;266:246–51.

Liu B, Furevi A, Perepelov A V. et al. Structure and genetics of Escherichia coli O antigens. FEMS Microbiol Rev 2020;44:655–83.

Liu B, Knirel YA, Feng L et al. Structural diversity in Salmonella O antigens and its genetic basis. FEMS Microbiol Rev 2014;38:56–89.

Liu H, Xiong Y, Liu X et al. Complete genome sequence of a novel virulent phage ST31 infecting Escherichia coli H21. Arch Virol 2018;163:1993–6.

Marin J, Clermont O, Royer G et al. The Population Genomics of Increased Virulence and Antibiotic Resistance in Human Commensal Escherichia coli over 30 Years in France. Appl Environ Microbiol 2022;88, DOI: 10.1128/aem.00664-22.

Nilsson E, Li K, Fridlund J et al. Genomic and Seasonal Variations among Aquatic Phages Infecting the Baltic Sea Gammaproteobacterium Rheinheimera sp. Strain BAL341. 2019, DOI: 10.1128/AEM.

Olszak T, Shneider MM, Latka A et al. The O-specific polysaccharide lyase from the phage LKA1 tailspike reduces Pseudomonas virulence. Sci Rep 2017;7:1–14.

Paterson DL, Bonomo RA. Extended-Spectrum Beta-Lactamases: a Clinical Update. Clin Microbiol Rev 2005;18:657–86.

Pires DP, Oliveira H, Melo LDR et al. Bacteriophage-encoded depolymerases: their diversity and biotechnological applications. Appl Microbiol Biotechnol 2016;100:2141–51.

Plattner M, Shneider MM, Arbatsky NP et al. Structure and Function of the Branched Receptor-Binding Complex of Bacteriophage CBA120. J Mol Biol 2019;431:3718–39.

Prokhorov NS, Riccio C, Zdorovenko EL et al. Function of bacteriophage G7C esterase tailspike in host cell adsorption. Mol Microbiol 2017;105:385–98.

Schmidt A, Rabsch W, Broeker NK et al. Bacteriophage tailspike protein based assay to monitor phase variable glucosylations in Salmonella O-antigens. BMC Microbiol 2016;16:1–11.

Scholl D, Cooley M, Williams SR et al. An engineered R-type pyocin is a highly specific and sensitive bactericidal agent for the food-borne pathogen Escherichia coli O157:H7. Antimicrob Agents Chemother 2009;53:3074–80.

Soffer N, Abuladze T, Woolston J et al. Bacteriophages safely reduce Salmonella contamination in pet food and raw pet food ingredients. Bacteriophage 2016;6:e1220347.

Sørensen AN, Woudstra C, Sørensen MCH et al. Subtypes of tail spike proteins predicts the host range of Ackermannviridae phages. Comput Struct Biotechnol J 2021;19, DOI: 10.1016/j.csbj.2021.08.030.

Steinbacher S, Baxa U, Miller S et al. Crystal structure of phage P22 tailspike protein complexed with Salmonella sp. O-antigen receptors. Proc Natl Acad Sci U S A 1996;93:10584–8.

Sullivan MJ, Petty NK, Beatson SA. Easyfig: A genome comparison visualizer. Bioinformatics 2011;27:1009–10.

Tamaki Y, Narimatsu H, Miyazato T et al. The Relationship between O-Antigens and Pathogenic Genes of Diarrhea-Associated Escherichia coli. Jpn J Infect Dis 2005a;58:65–9.

Tamaki Y, Narimatsu H, Miyazato T et al. The relationship between O-antigens and pathogenic genes of diarrhea-associated Escherichia coli. Jpn J Infect Dis 2005b;58:65–69.

Thanh NC, Nagayoshi Y, Fujino Y et al. Characterization and Genome Structure of Virulent Phage EspM4VN to Control Enterobacter sp. M4 Isolated From Plant Soft Rot. Front Microbiol 2020;11, DOI: 10.3389/fmicb.2020.00885.

Turner D, Kropinski AM, Adriaenssens EM. A roadmap for genome-based phage taxonomy. Viruses 2021;13, DOI: 10.3390/v13030506.

Vitt AR, Sørensen AN, Bojer MS et al. A collection of diverse bacteriophages for biocontrol of ESBL- and AmpC-β-lactamase-producing E. coli. bioRxiv 2023:2023.09.14.557699.

Walter M, Fiedler C, Grassl R et al. Structure of the Receptor-Binding Protein of Bacteriophage Det7: a Podoviral Tail Spike in a Myovirus. J Virol 2008;82:2265–73.

Wang H, Liu Y, Bai C et al. Translating bacteriophage-derived depolymerases into antibacterial therapeutics: Challenges and prospects. Acta Pharm Sin B 2023, DOI: 10.1016/j.apsb.2023.08.017.

Waseh S, Hanifi-Moghaddam P, Coleman R et al. Orally administered P22 phage tailspike protein reduces Salmonella colonization in chickens: Prospects of a novel therapy against bacterial infections. PLoS One 2010;5, DOI: 10.1371/journal.pone.0013904.

Williams SR, Gebhart D, Martin DW et al. Retargeting R-type pyocins to generate novel bactericidal protein complexes. Appl Environ Microbiol 2008;74:3868–76.

Witte S, Zinsli L V., Gonzalez-Serrano R et al. Structural and functional characterization of the receptor binding proteins of Escherichia coli O157 phages EP75 and EP335. Comput Struct Biotechnol J 2021;19:3416–26.

Yan T, Liang L, Yin P et al. Application of a novel phage LPST94 for biological control of Salmonella in foods. Microorganisms 2020;8, DOI: 10.3390/microorganisms8030400.

Zampara A, Sørensen MCH, Gencay YE et al. Developing Innolysins Against Campylobacter jejuni Using a Novel Prophage Receptor-Binding Protein. Front Microbiol 2021;12, DOI: 10.3389/fmicb.2021.619028.

Zampara A, Sørensen MCH, Grimon D et al. Exploiting phage receptor binding proteins to enable endolysins to kill Gram-negative bacteria. Sci Rep 2020;10, DOI: 10.1038/s41598-020-68983-3.

