## Supplementary information for "Agtrevirus phage AV101 infect diverse extended spectrum β-lactamase *E. coli* by recognizing four different O-antigens"

### Supplementary data

**Table S1: A list of all phage genomes used for in-silico analysis and their identity to phage AV101.**

| Genus* | Phages | Query cover | Percent identity | Accession |
|---|---|---|---|---|
| **Unclassified Aglimvirinae** | Enterobacter phage fGh-Ecl02 | 87% | 97.54% | ON212266.1 |
| | Escherichia phage PH4 | 85% | 97.83% | ON184126.1 |
| | Escherichia phage PC3 | 85% | 97.83% | ON184125.1 |
| | Escherichia phage vB_EcoM-RPN242 | 88% | 97.74% | OL656110.1 |
| | Escherichia phage vB_EcoM-ZQ1 | 85% | 97.57% | MW650886.1 |
| **Agtrevirus** | Shigella phage phiSboM-AG3 | 85% | 92.25% | NC_013693.1 |
| | Salmonella phage P46FS4 | 85% | 92.08% | NC_049509.1 |
| | Salmonella phage SKML-39 | 85% | 91.76% | JX181829.1 |
| | Shigella phage vB_SboS_Gloob | 85% | 92% | OL615011.1 |
| | Shigella phage vB_SboM_ChubbyThor | 85% | 93.03% | OL615013.1 |
| | Shigella phage MK-13 | 84% | 91.94% | NC_049455.1 |
| | Enterobacter phage EspM4VN | 83% | 90.23% | NC_049384.1 |
| | Salmonella phage pSal-SNUABM-02 | 82% | 90.52% | MT857002.1 |

**Table S2: overview of genes marked in Figure 1.** The genes that are highlighted in Figure 1 are listed below in the order seen in the figure (right to left).

| Phage SNUAMB-02 | Gene product | Function | Position | Colour on Fig 1 |
|---|---|---|---|---|
| | 105 | hypothetical protein | 2620..3789 | Blue |
| | 106 | Putative tail spike protein | 3842..6877 | Blue |
| | 107 | Putative tail spike protein | 6930..9353 | Blue |
| | 128 | hypothetical protein | 29730..31052 | Orange |
| | 202 | putative ribonucleoside-diphosphate reductase subunit alpha | 79811..80470 | Red |
| | 203 | putative ribonucleotide-diphosphate reductase<br>subunit beta | 80638..82812 | Red |
| | 5 | Putative thymidylate synthase | 97785..98357 | Red |
| | 6 | putative dNMP kinase | 98374..99423 | Red |
| | 7 | putative dCTP pyrophosphatase | 99423..99989 | Red |
| | 84 | putative homing endonuclease | 144086..144283 | Orange |
| Phage EspM4VN | Gene product | Function | Position | Colour on Fig. 1 |
| | 213 | hypothetical protein | 2620..3831 | Blue |
| | 214 | hypothetical protein | 3884..7213 | Blue |
| | 215 | hypothetical protein | 7259..9055 | Blue |
| | 216 | hypothetical protein | 9110..11674 | Blue |
| | 165 | ribose-phosphate pyrophosphokinase | 50312..51106 | Green |
| | 164 | nicotinamide phosphoribosyltransferase | 51217..52071 | Green |
| | 119 | ribonucleoside-diphosphate reductase | 80789..81631 | Red |
| | 118 | NrdB ribonucleotide reductase subunit beta | 81717..83996 | Red |
| | 97 | part of thymidylate synthase | 97641..98027 | Red |
| | 96 | part of thymidylate synthase | 98268..98534 | Red |
| | 95 | part of thymidylate synthase | 98492..98833 | Red |
| | 94 | hypothetical protein | 98817..98978 | Red |
| | 1 | putative dUTP diphosphatase | 99231..99611 | Red |

| | | | | |
|---|---|---|---|---|
| | 76 | hypothetical protein | 143742..143939 | Orange |
| Phage fGh-Ecl02 | Protein Id | Function | Position | Colour on Fig 1 |
| | USL85809.1 | hypothetical protein | 3857..6853 | Blue |
| | USL85810.1 | hypothetical protein | 6932..9034 | Blue |
| | USL85811.1 | hypothetical protein | 9114..11777 | Blue |
| | USL85812.1 | hypothetical protein | 11835..14468 | Blue |
| | USL85832.1 | putative homing endonuclease | 34301..35827 | Orange |
| | USL85896.1 | ribonucleoside-diphosphate reductase subunit alpha | 79932..82214 | Red |
| | USL85897.1 | ribonucleotide reductase beta subunit | 82285..83388 | Red |
| | USL85921.1 | putative dUTP diphosphatase | 97956..98510 | Red |
| | USL85922.1 | hypothetical protein | 98510..99055 | Red |
| | USL85923.1 | RecA-like recombination protein | 99040..100125 | Red |
| | USL85792.1 | putative homing endonuclease | 138315..139913 | Orange |
| Phage AV101 | Gene product | Function | Position | Colour on Fig x |
| | Peg 193 | Tail spike protein 4 | 3884..7162 | Blue |
| | Peg 192 | Tail spike protein 3 | 7466..9412 | Blue |
| | Peg 191 | Hypothetical protein | 9445..9600 | Blue |
| | Peg 190 | Tail spike protein 2 | 9952..12240 | Blue |
| | Peg 189 | Tail spike protein 1 | 12288..14489 | Blue |
| | Peg 188 | Hypothetical protein | 14832..14957 | Blue |
| | Peg 169 | hypothetical protein | 34502..35974 | Orange |
| | Peg 141 | Ribose-phosphate pyrophosphokinase | 53586..54440 | Green |
| | Peg 140 | Nicotinamide phosphoribosyltransferase | 54437..56101 | Green |
| | Peg 101 | ribonucleotide reductase of class Ia (aerobic) alpha subunit | 83581..85863 | Red |
| | Peg 100 | ribonucleotide reductase of class Ia (aerobic) beta subunit | 85934..87037 | Red |
| | Peg 78 | Putative thymidylate synthase | 99997..101043 | Red |
| | Peg 77 | Hypothetical protein | 101040..101609 | Red |
| | Peg 76 | putative dUTP diphosphatase | 101606..102160 | Red |
| | Peg 14 | putative homing endonuclease | 141564..142481 | Orange |

| | | | | |
|---|---|---|---|---|
| | Peg 13 | putative homing endonuclease | 142474..143160 | Orange |
| Phage ZQ1 | Protein id | Function | Position | Colour on Fig. 1 |
| | QVW27043.1 | Tail fibers protein | 3884..7006 | Blue |
| | QVW27042.1 | Tail spike protein | 7106..9001 | Blue |
| | QVW27222.1 | Hypothetical protein | 9221..9685 | Blue |
| | QVW27221.1 | hypothetical protein | 9756..10157 | Blue |
| | QVW27209.1 | Putative homing endonuclease | 22741..24525 | Orange |
| | QVW27201.1 | Putative homing endonuclease | 30520..31992 | Orange |
| | QVW27144.1 | ribonucleotide reductase of class Ia (aerobic) alpha subunit | 75315..77597 | Red |
| | QVW27143.1 | ribonucleotide reductase of class Ia (aerobic) beta subunit | 77668..78771 | Red |
| | QVW27122.1 | Putative thymidylate synthase | 91465..92511 | Red |
| | QVW27121.1 | putative deoxynucleotide monophosphate kinase | 92508..93077 | Red |
| | QVW27120.1 | putative dUTP diphosphatase | 93074..93628 | Red |
| | QVW27062.1 | putative homing endonuclease | 133656..135254 | Orange |
| Phage PC3 | Gene product | Function | Position | Colour on Fig. 1 |
| | 1 | hypothetical protein | 3875..7036 | Blue |
| | 2 | Tail spike protein 3 | 7136..9685 | Blue |
| | 3 | Tail spike protein 2 | 9961..12384 | Blue |
| | 4 | Tail spike protein | 12703..14928 | Blue |
| | 16 | homing endonuclease | 28584..30272 | Orange |
| | 25 | homing endonuclease | 36488..38353 | Orange |
| | 88 | ribonucleotide reductase of class Ia (aerobic) alpha subunit | 81541..83823 | Red |
| | 89 | Hypothetical protein | 83893..84186 | Red |
| | 90 | ribonucleotide reductase of class Ia (aerobic) beta subunit | 84188..85291 | Red |
| | 112 | thymidylate synthase | 98260..99306 | Red |
| | 113 | hypothetical protein | 99303..99872 | Red |
| | 114 | putative dUTP diphosphatase | 99869..100423 | Red |
| Phage PH4 | Gene product | Function | Position | Colour on Fig. 1 |
| | 1 | hypothetical protein | 3875..7036 | Blue |
| | 2 | Tail spike protein 3 | 7136..9685 | Blue |

| | | | | |
|---|---|---|---|---|
| | 3 | Tail spike protein 2 | 9961..12384 | Blue |
| | 4 | Tail spike protein | 12703..14928 | Blue |
| | 16 | homing endonuclease | 28584..30272 | Orange |
| | 25 | homing endonuclease | 36488..38353 | Orange |
| | 88 | ribonucleotide reductase of class Ia (aerobic) alpha subunit | 81541..83823 | Red |
| | 89 | Hypothetical protein | 83893..84186 | Red |
| | 90 | ribonucleotide reductase of class Ia (aerobic) beta subunit | 84188..85291 | Red |
| | 112 | thymidylate synthase | 98260..99306 | Red |
| | 113 | hypothetical protein | 99303..99872 | Red |
| | 114 | putative dUTP diphosphatase | 99869..100423 | Red |
| Phage RPN242 | Gene product | Function | Position | Colour on Fig x |
| | 164 | Tail fibers protein | 2620..3810 | Blue |
| | 163 | Tail fiber protein | 3863..7024 | Blue |
| | 162 | Tail spike protein | 7124..9673 | Blue |
| | 161 | hypothetical protein | 9792..12200 | Blue |
| | 142 | homing endonuclease | 33187..34509 | Orange |
| | 76 | ribonucleoside-diphosphate reductase subunit alpha | 79585..80427 | Red |
| | 75 | Hypothetical protein | 80511..82793 | Red |
| | 74 | ribonucleotide-diphosphate reductase subunit beta | 82863..83156 | Red |
| | 52 | thymidylate synthase | 96571..97230 | Red |
| | 51 | Hypothetical protein | 97230..98276 | |
| | 50 | 2-deoxyuridine 5-triphosphate nucleotidohydrolase | 98273..98842 | Red |
| | 184 | putative homing endonuclease | 138680..138877 | Orange |
| | 183 | homing endonuclease | 138880..139785 | Orange |
| Phage SKML-39 | Gene product | Function | Position | Colour on Fig x |
| | 207 | hypothetical protein | 3884..6943 | Blue |
| | 208 | hypothetical protein | 6991..8916 | Blue |
| | 1 | Hypothetical protein | 9543..11813 | Blue |
| | 2 | hypothetical protein | 11863..14001 | Blue |

| | | | | |
|---|---|---|---|---|
| | 48 | ribose-phosphate pyrophosphokinase | 52051..52905 | Green |
| | 49 | nicotinamide phosphoribosyltransferase | 52902..54566 | Green |
| | 90 | ribonucleotide reductase of class Ia (aerobic) alpha subunit | 81930..84212 | Red |
| | 91 | Hypothetical protein | 84283..84576 | |
| | 92 | ribonucleotide reductase of class Ia (aerobic) beta subunit | 84578..85681 | Red |
| | 115 | Hypothetical protein | 99516..100562 | Red |
| | 116 | Hypothetical protein | 100562..101128 | Red |
| | 117 | Hypothetical protein | 101125..101679 | Red |
| | 186 | Hypothetical protein | 143031..143948 | Orange |
| Phage Gloob | Gene product | Function | Position | Colour on Fig. 1 |
| | 169 | putative right handed beta helix region family | 2635..3846 | Blue |
| | 168 | Putative tail spike protein | 3869..6901 | Blue |
| | 167 | Putative tail spike protein | 6931..8979 | Blue |
| | 166 | putative right-handed parallel beta-helix | 9164..11386 | Blue |
| | 121 | putative ribose-phosphate pyrophosphokinase | 49301..49540 | Green |
| | 120 | putative nicotinamide | 49801..50655 | Green |
| | 79 | putative ribonucleoside-diphosphate | 79633..80508 | Red |
| | 78 | hypothetical protein | 80594..82873 | Red |
| | 77 | putative ribonucleotide reductase subunit | 82944..83237 | Red |
| | 55 | putative thymidylate synthase | 97367..98413 | Red |
| | 54 | putative P-loop containing nuceloside | 97367..98413 | Red |
| | 53 | putative dUTP diphosphatase | 98410..98979 | Red |
| | 195 | putative homing endonuclease | 141345..141548 | Orange |
| Phage ChubbyThor | Gene product | Function | Position | Colour on Fig. 1 |

| | | | | |
|---|---|---|---|---|
| | 3 | putative right handed beta helix region family | 2635..3846 | Blue |
| | 2 | putative head binding protein | 3869..6793 | Blue |
| | 1 | putative tail fibers protein | 6875..9007 | Blue |
| | 208 | putative tail protein | 9138..11843 | Blue |
| | 189 | Putative HNH endonuclease | 31758..33089 | Orange |
| | 163 | putative ribose-phosphate pyrophosphokinase | 51049..51843 | Green |
| | 162 | putative nicotinamide | 51954..52808 | Green |
| | 118 | putative ribonucleotide diphosphate reductase | 82796..83455 | Red |
| | 117 | putative ribonucleotide-diphosphate reductase | 83515..85803 | Red |
| | 95 | Putative thymidylate synthase | 99046..99720 | Red |
| | 94 | putative dNMP kinase | 99717..100766 | Red |
| | 93 | putative dUTP diphosphatase | 100763..101332 | Red |
| | 27 | Hypothetical proteins | 143610..143813 | Orange |
| | 26 | putative endonuclease | 143810..144727 | Orange |
| Phage Ag3 | Gene product | Function | Position | Colour on Fig. 1 |
| | 178 | hypothetical protein | 2620..3831 | Blue |
| | 177 | Tailspike protein | 3884..6943 | Blue |
| | 176 | Tailspike protein | 6990..9011 | Blue |
| | 175 | hypothetical protein | 9212..11473 | Blue |
| | 130 | putative nicotinamide phosphoribosyl transferase | 49705..51348 | Green |
| | 83 | NrdA ribonucleoside-diphosphate reductase alpha subunit | 80026..82305 | Red |
| | 82 | Hypothetical protein | 82376..82669 | Red |
| | 81 | NrdB ribonucleotide reductase subunit beta | 82671..83774 | Red |
| | 59 | Putative thymidylate synthase | 96118..96780 | Red |
| | 58 | Hypothetical protein | 96780..97826 | Red |
| | 57 | putative dUTP diphosphatase | 97823..98392 | Red |
| | 201 | Hypothetical protein | 141295..141492 | Orange |
| | 200 | SegD-like protein | 141495..142412 | Orange |

Supplementary data - Agtrevirus phage AV101 infect diverse extended spectrum β-lactamase E. coli by recognizing four different O-antigens

| Phage P46FS4 | Gene product | Function | Position | Colour on Fig. |
|---|---|---|---|---|
| | 1 | hypothetical protein | 3884..6943 | Blue |
| | 2 | Tail spike protein | 6990..7469 | Blue |
| | 3 | Hypothetical protein | 7450..8997 | Blue |
| | 4 | Tail spike protein | 9053..11476 | Blue |
| | 5 | Tail fibers protein | 11639..14101 | Blue |
| | 50 | ribose-phosphate pyrophosphokinase | 50468..51322 | Green |
| | 51 | nicotinamide phosphoribosyltransferase | 51319..52983 | Green |
| | 94 | ribonucleotide reductase of class Ia (aerobic) alpha subunit | 81226..83505 | Red |
| | 95 | Hypothetical protein | 83576..83869 | Red |
| | 96 | ribonucleotide reductase of class Ia (aerobic) beta subunit | 83871..84974 | Red |
| | 118 | Putative thymidylate synthase | 97995..99041 | Red |
| | 119 | Hypothetical protein | 99038..99607 | Red |
| | 120 | putative dUTP diphosphatase | 99604..100158 | Red |
| | 190 | SegD-like protein | 142956..143642 | Orange |
| Phage MK-13 | Gene product | Function | Position | Colour on Fig. 1 |
| | 202 | hypothetical protein | 2620..3831 | Blue |
| | 201 | hypothetical protein | 3884..6778 | Blue |
| | 200 | Hypothetical protein | 6860..8962 | Blue |
| | 199 | hypothetical protein | 9040..11820 | Blue |
| | 154 | hypothetical protein | 49429..50223 | Green |
| | 153 | hypothetical protein | 50334..51188 | Green |
| | 110 | ribonucleoside-diphosphate reductase large subunit | 80661..81503 | Red |
| | 109 | hypothetical protein | 81589..83868 | Red |
| | 6 | hypothetical protein | 98258..98920 | Red |
| | 7 | hypothetical protein | 98920..99966 | Red |
| | 8 | hypothetical protein | 99963..100532 | Red |
| | 75 | hypothetical protein | 141787..141984 | Orange |

**Table S3: In-silico analysis of TSP1-4.** TSP1 and TSP3 show sequence similarity to *E. coli* genomes suggesting similarity to prophages. TSP2 and TSP4 show similarity to other phages.

| | **Hit** | **Organism** | **Accession number genome** | **Query cover** | **Percentage identity** | **Protein ID** |
|---|---|---|---|---|---|---|
| **TSP1** | | | | | | |
| | Phage head-binding domain-containing protein | Escherichia coli H001 contig 1.26 | ADIW01000026 | 78% | 66.49% | OSK75853.1 |
| | Phage tailspike protein | Escherichia coli strain 89 contig 15 | JAIZTL010000015 | 78% | 66.32% | MCA8532455.1 |
| | Phage head-binding domain-containing protein | Escherichia coli CHS 77 contig 1.1.C4 | JMVO01000004 | 78% | 65.97% | KDG58944.1 |
| **TSP2** | Hypothetical protein | *E. coli phage A221* | ON862890.1 | 100% | 100% | UTQ72550.1 |
| | Hypothetical protein | Escherichia coli isolate CP21 | DADPIP010000000 | 68% | 91.73% | HAZ7467268.1 |
| | Tail spike protein | Salmonella phage UAB_1 | OL656106.1 | 68% | 92.12% | UIS31512.1 |
| **TSP3** | | | | | | |
| | Hypothetical protein | Escherichia coli strain 705003 MRSN705003_contig00003 | JAKYFG010000003 | 85% | 83.63% | MCN8021000.1 |
| | Hypothetical protein | Escherichia coli strain 767377 MRSN767377_contig00056 | JAMEID010000056 | 85% | 83.63% | MCN3003243.1 |
| | Hypothetical protein | Escherichia coli strain VRES0635 | UDHQ01000018 | 85% | 83.45% | MCY6644629.1 |
| **TSP4** | | | | | | |
| | Tail fiber protein | phage SalM-LPST94 | MH523359.1 | 68% | 90.91% | AXF41694.1 |
| | Tail spike protein | phage vB_EcoM-Ro121c4YLVW | MH051333.1 | 61% | 70.49% | YP_009984889.1 |
| | Tail fiber protein | phage ST31 | KY962008.1 | 60% | 68.86% | YP_009790654.1 |

.

**Table S4: Prophage identified in the *E. coli* sequences.** PHASTER analysis of the *E. coli* genomes that carried proteins with similarity towards the TSPs of AV101.

| | **Assesion number** | **Prophage length** | **Completeness (Score)** | **No. of proteins** | **Region position** | **Most common phage** | **GC %** |
|---|---|---|---|---|---|---|---|
| **TSP1:** | | | | | | | |
| Escherichia coli H001 contig 1.26 | ADIW01000026 | 27 Kb | Questionable (70) | 33 | 413-27507 | PHAGE_Entero_Sf101 | 46.03 |
| Escherichia coli strain 89 contig 15 | JAIZTL010000015 | 44.2 Kb | Intact (150) | 60 | 39194-83481 | PHAGE_Entero_Sf101 | 47.85 |
| Escherichia coli CHS 77 contig 1.1.C4 | JMVO01000004 | 41.9 Kb | Intact (120) | 57 | 25895-67861 | PHAGE_Entero_IME10 | 47.29 |
| **TSP2:** | | | | | | | |
| Escherichia coli isolate CP21 | DADPIP010000000 | 145.9 Kb | Intact (133) | 236 | 326-146309 | PHAGE_Entero_phi92 | 37.48 |
| **TSP3:** | | | | | | | |
| Escherichia coli strain 705003 MRSN705003_contig00003 | JAKYFG010000003 | 43.6 Kb | Intact (150) | 60 | 280671-324348 | PHAGE_Entero_mEp390 | 47.16 |
| Escherichia coli strain 767377 MRSN767377_contig00056 | JAMEID010000056 | 43.7 Kb | Intact (150) | 61 | 19221-62927 | PHAGE_Entero_mEp390 | 47.17 |
| Escherichia coli strain VRES0635, contig: ERS792374SCcontig000018 | UDHQ01000018 | 14.1 Kb | Questionable (80) | 21 | 11789-25952 | PHAGE_Entero_HK140 | 44.15 |

#### Table S5: List of bacteria used in the study.

| Genus | Species or serovar | KU ID | Serotype | Origin | Rerence |
|---|---|---|---|---|---|
| *Escherichia* | coli | BL21(DE3) | ND | Invitrogen | Protein expression |
| *Escherichia* | coli | Stellar cells | ND | Takara Bio | Plasmid propagation strain |
| *Salmonella* | Typhimurium | LT2c | O4 | (54) | Host range analysis |
| *Salmonella* | Derby | EGS1 | O4 | JEO | Host range analysis |
| *Salmonella* | Enteritidis | EGS48 | O9 | JEO | Host range analysis |
| *Salmonella* | Seftenberg | EGS381 | O3 | JEO | Host range analysis |
| *Salmonella* | Anatum | EGS388 | O3,10 | JEO | Host range analysis |
| *Salmonella* | Onderstepoort | EGS391 | O6 | JEO | Host range analysis |
| *Salmonella* | Minnesota | JEO3 | O21 | JEO | Host range analysis |
| *Escherichia* | coli | ESBL038 | O8 | In prep. | Host range analysis |
| *Escherichia* | coli | ESBL040 | O8 | In prep. | Host range analysis |
| *Escherichia* | coli | ESBL048 | O82 | In prep. | Host range analysis |
| *Escherichia* | coli | ESBL079 | O82 | In prep. | Host range analysis |
| *Escherichia* | coli | ESBL082 | O82 | In prep. | Host range analysis |
| *Escherichia* | coli | ESBL058 | O153 | In prep. | Host range analysis |
| *Escherichia* | coli | ESBL078 | O153 | In prep. | Host range analysis |
| *Escherichia* | coli | ESBL089 | O153 | In prep. | Host range analysis |
| *Escherichia* | coli | ESBL144 | O159 | In prep. | Host range analysis |
| *Escherichia* | coli | ECOR1 | O144:H4 | (57) | Host range analysis |
| *Escherichia* | coli | ECOR2 | O48:H32 | (57) | Host range analysis |
| *Escherichia* | coli | ECOR3 | O1:H32 | (57) | Host range analysis |
| *Escherichia* | coli | ECOR4 | OR:H? | (57) | Host range analysis |
| *Escherichia* | coli | ECOR5 | O?:H6 | (57) | Host range analysis |
| *Escherichia* | coli | ECOR6 | O173:H? | (57) | Host range analysis |
| *Escherichia* | coli | ECOR7 | O8:H45 | (57) | Host range analysis |
| *Escherichia* | coli | ECOR8 | O86:H2 | (57) | Host range analysis |
| *Escherichia* | coli | ECOR9 | O167:H- | (57) | Host range analysis |
| *Escherichia* | coli | ECOR10 | O6:H10 | (57) | Host range analysis |
| *Escherichia* | coli | ECOR11 | O10:H- | (57) | Host range analysis |
| *Escherichia* | coli | ECOR12 | O?:H32 | (57) | Host range analysis |
| *Escherichia* | coli | ECOR13 | OR:H25 | (57) | Host range analysis |
| *Escherichia* | coli | ECOR14 | O71:H4 | (57) | Host range analysis |
| *Escherichia* | coli | ECOR15 | O25:H30 | (57) | Host range analysis |
| *Escherichia* | coli | ECOR16 | O9:H10 | (57) | Host range analysis |

| | | | | | |
|---|---|---|---|---|---|
| *Escherichia* | coli | ECOR17 | O29:H- | (57) | Host range analysis |
| *Escherichia* | coli | ECOR18 | O?:H11 | (57) | Host range analysis |
| *Escherichia* | coli | ECOR19 | O89:H? | (57) | Host range analysis |
| *Escherichia* | coli | ECOR20 | O121:H11 | (57) | Host range analysis |
| *Escherichia* | coli | ECOR21 | O121:H11 | (57) | Host range analysis |
| *Escherichia* | coli | ECOR22 | O150:H28 | (57) | Host range analysis |
| *Escherichia* | coli | ECOR23 | O25.H1 | (57) | Host range analysis |
| *Escherichia* | coli | ECOR24 | O15:H- | (57) | Host range analysis |
| *Escherichia* | coli | ECOR25 | O127:H40 | (57) | Host range analysis |
| *Escherichia* | coli | ECOR26 | O104:H21 | (57) | Host range analysis |
| *Escherichia* | coli | ECOR27 | O104:H21 | (57) | Host range analysis |
| *Escherichia* | coli | ECOR28 | O104:H21 | (57) | Host range analysis |
| *Escherichia* | coli | ECOR29 | O150:H21 | (57) | Host range analysis |
| *Escherichia* | coli | ECOR30 | O113:H21 | (57) | Host range analysis |
| *Escherichia* | coli | ECOR31 | O79:H25 | (57) | Host range analysis |
| *Escherichia* | coli | ECOR32 | O25:H1 | (57) | Host range analysis |
| *Escherichia* | coli | ECOR33 | O7:H21 | (57) | Host range analysis |
| *Escherichia* | coli | ECOR34 | O88:H- | (57) | Host range analysis |
| *Escherichia* | coli | ECOR35 | O1:H- | (57) | Host range analysis |
| *Escherichia* | coli | ECOR36 | O1:H- | (57) | Host range analysis |
| *Escherichia* | coli | ECOR37 | O55:H7 | (57) | Host range analysis |
| *Escherichia* | coli | ECOR38 | O7:H- | (57) | Host range analysis |
| *Escherichia* | coli | ECOR39 | O7:H- | (57) | Host range analysis |
| *Escherichia* | coli | ECOR40 | O7:H- | (57) | Host range analysis |
| *Escherichia* | coli | ECOR41 | O7:H- | (57) | Host range analysis |
| *Escherichia* | coli | ECOR42 | O87:H26 | (57) | Host range analysis |
| *Escherichia* | coli | ECOR43 | O?:H18 | (57) | Host range analysis |
| *Escherichia* | coli | ECOR44 | O17:H34 | (57) | Host range analysis |
| *Escherichia* | coli | ECOR45 | O?:H2 | (57) | Host range analysis |
| *Escherichia* | coli | ECOR46 | O1:H- | (57) | Host range analysis |
| *Escherichia* | coli | ECOR47 | O17:H18 | (57) | Host range analysis |
| *Escherichia* | coli | ECOR48 | O23:H15 | (57) | Host range analysis |
| *Escherichia* | coli | ECOR49 | O2:H4 | (57) | Host range analysis |
| *Escherichia* | coli | ECOR50 | O2:H4 | (57) | Host range analysis |
| *Escherichia* | coli | ECOR51 | O25:H1 | (57) | Host range analysis |
| *Escherichia* | coli | ECOR52 | O25:H1 | (57) | Host range analysis |

| | | | | | |
|---|---|---|---|---|---|
| *Escherichia* | coli | ECOR53 | O4:H5 | (57) | Host range analysis |
| *Escherichia* | coli | ECOR54 | O25:H1 | (57) | Host range analysis |
| *Escherichia* | coli | ECOR55 | O25:H1 | (57) | Host range analysis |
| *Escherichia* | coli | ECOR56 | O6:H10 | (57) | Host range analysis |
| *Escherichia* | coli | ECOR57 | O2:H1 | (57) | Host range analysis |
| *Escherichia* | coli | ECOR58 | O112:H8 | (57) | Host range analysis |
| *Escherichia* | coli | ECOR59 | O2:H4 | (57) | Host range analysis |
| *Escherichia* | coli | ECOR60 | O4:H5 | (57) | Host range analysis |
| *Escherichia* | coli | ECOR61 | O2:H4 | (57) | Host range analysis |
| *Escherichia* | coli | ECOR62 | O2:H4 | (57) | Host range analysis |
| *Escherichia* | coli | ECOR63 | OR:H- | (57) | Host range analysis |
| *Escherichia* | coli | ECOR64 | O75:H- | (57) | Host range analysis |
| *Escherichia* | coli | ECOR65 | O8:H10 | (57) | Host range analysis |
| *Escherichia* | coli | ECOR66 | O4:H40 | (57) | Host range analysis |
| *Escherichia* | coli | ECOR67 | O141:H49 | (57) | Host range analysis |
| *Escherichia* | coli | ECOR68 | O25:H21 | (57) | Host range analysis |
| *Escherichia* | coli | ECOR69 | O86:H10 | (57) | Host range analysis |
| *Escherichia* | coli | ECOR70 | O78:H- | (57) | Host range analysis |
| *Escherichia* | coli | ECOR71 | OR:H19 | (57) | Host range analysis |
| *Escherichia* | coli | ECOR72 | O8:H30 | (57) | Host range analysis |
| *Escherichia* | coli | NCTC12900 | O157:H7 | JEO | Host range analysis |
| *Escherichia* | coli | ATCC35150 | O157:H7 | JEO | Host range analysis |
| *Escherichia* | coli | ATCC43888 | O157:H7 | JEO | Host range analysis |
| *Escherichia* | coli | ATCC43895 | O157:H7 | JEO | Host range analysis |
| *Escherichia* | coli | JEO1894 | O149:K88 | JEO | Host range analysis |
| *Escherichia* | coli | MG1665 | K12 | | Host range analysis |

### Figure S1

| | | 1 | 2 | 3 | 4 | 5 | 6 | 7 | 8 | 9 | 10 | 11 | 12 | 13 | 14 |
|---|---|---|---|---|---|---|---|---|---|---|---|---|---|---|---|
| Salmonella phage pSal-SNUABM-02 | 1 | | 86,64 | 87,58 | 87,09 | 87,28 | 86,98 | 88,10 | 87,86 | 87,69 | 88,05 | 86,89 | 87,06 | 88,15 | 87,09 |
| Enterobacter phage EspM4VN | 2 | 86,64 | | 88,79 | 87,67 | 87,76 | 87,65 | 89,40 | 88,95 | 88,68 | 89,18 | 87,81 | 87,86 | 88,89 | 87,67 |
| Shigella phage vB_SboM_ChubbyThor | 3 | 87,58 | 88,79 | | 90,53 | 89,84 | 90,93 | 95,47 | 96,59 | 94,78 | 95,97 | 90,35 | 90,32 | 95,07 | 90,52 |
| Escherichia phage PH4 | 4 | 87,09 | 87,67 | 90,53 | | 96,82 | 97,24 | 90,87 | 90,80 | 90,61 | 90,91 | 96,77 | 95,85 | 90,55 | 100,00 |
| Escherichia phage vB_EcoM-ZQ1 | 5 | 87,28 | 87,76 | 89,84 | 96,82 | | 96,75 | 90,97 | 90,62 | 90,23 | 90,79 | 95,86 | 96,16 | 90,20 | 96,82 |
| Escherichia phage vB_EcoM-RPN242 | 6 | 86,98 | 87,65 | 90,93 | 97,24 | 96,75 | | 90,97 | 91,11 | 90,93 | 91,39 | 96,13 | 96,00 | 90,90 | 97,24 |
| Salmonella phage P46FS4 | 7 | 88,10 | 89,40 | 95,47 | 90,87 | 90,97 | 90,97 | | 95,91 | 95,19 | 95,93 | 90,76 | 91,02 | 95,51 | 90,87 |
| Shigella phage Ag3 | 8 | 87,86 | 88,95 | 96,59 | 90,80 | 90,62 | 91,11 | 95,91 | | 95,34 | 95,99 | 90,56 | 90,57 | 95,44 | 90,80 |
| Salmonella phage SKML-39 | 9 | 87,69 | 88,68 | 94,78 | 90,61 | 90,23 | 90,93 | 95,19 | 95,34 | | 95,58 | 90,01 | 90,39 | 95,46 | 90,61 |
| Shigella phage MK-13 | 10 | 88,05 | 89,18 | 95,97 | 90,91 | 90,79 | 91,39 | 95,93 | 95,99 | 95,58 | | 90,86 | 90,97 | 95,41 | 90,91 |
| Enterobacter phage fGh-Ecl02 | 11 | 86,89 | 87,81 | 90,35 | 96,77 | 95,86 | 96,13 | 90,76 | 90,56 | 90,01 | 90,86 | | 96,75 | 90,25 | 96,77 |
| Escherichia phage AV101 | 12 | 87,06 | 87,86 | 90,32 | 95,85 | 96,16 | 96,00 | 91,02 | 90,57 | 90,39 | 90,97 | 96,75 | | 90,49 | 95,84 |
| Shigella phage vB_SboS_Gloob | 13 | 88,15 | 88,89 | 95,07 | 90,55 | 90,20 | 90,90 | 95,51 | 95,44 | 95,46 | 95,41 | 90,25 | 90,49 | | 90,55 |
| Escherichia phage PC3 | 14 | 87,09 | 87,67 | 90,52 | 100,00 | 96,82 | 97,24 | 90,87 | 90,80 | 90,61 | 90,91 | 96,77 | 95,84 | 90,55 | |

**Figure S1: Average Nucleotide Identity of the 14 Agtrevirus genomes.** All 14 phages were compared and showed that they all belong to the Agtrevirus genus.

### Figure S2

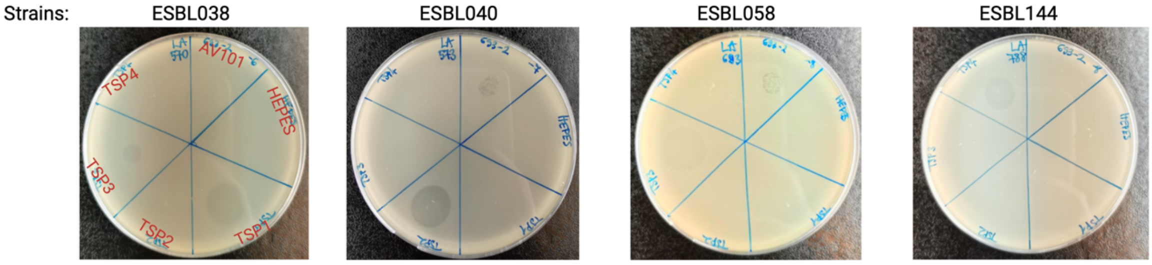


**Figure S2: Each TSPs of AV101 recognize a different AV101 hosts**. Each purified TSPs and AV101 were spotted on the known AV101 phage hosts to investigate their receptor binding ability. The protein buffer (HEPES) was added as a negative control.

### Figure S3

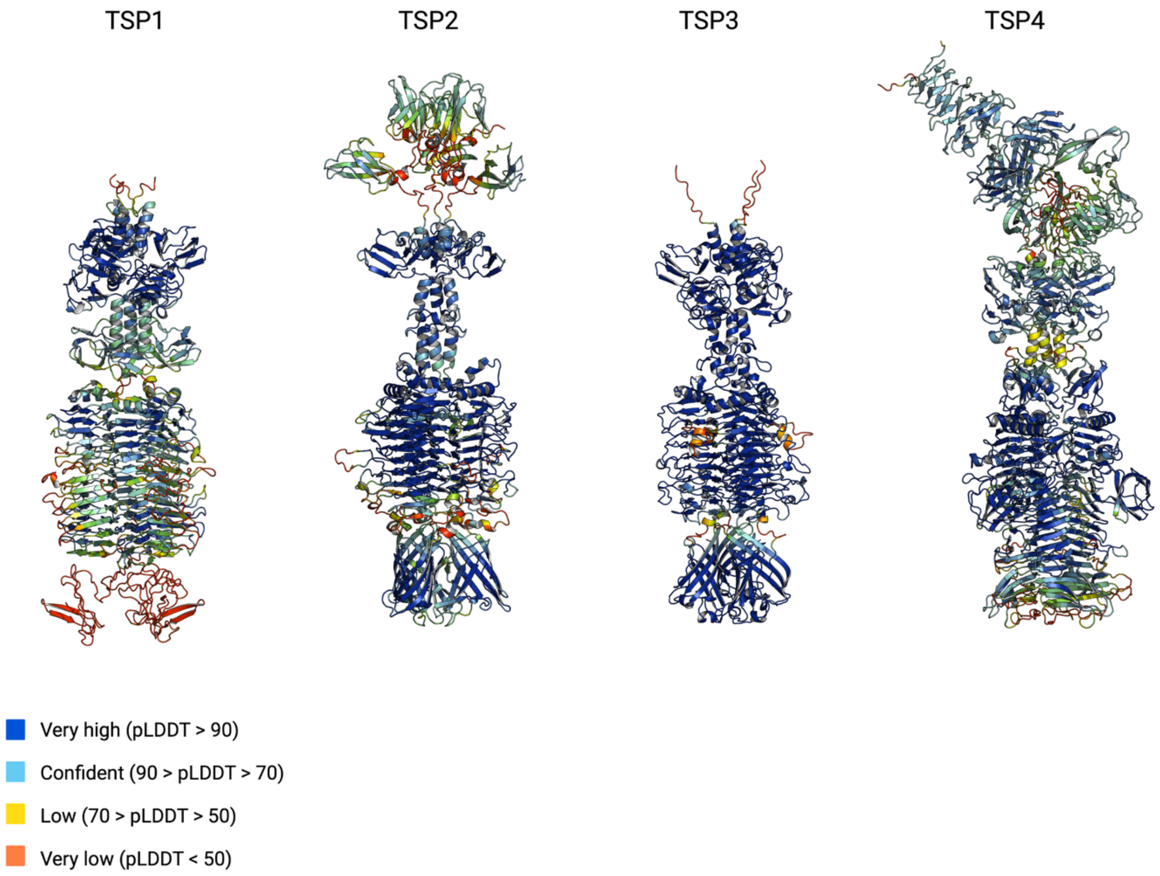


**Figure S3: AlphaFold2 prediction of the four AV101 TSPs.** The structures of the TSPs of AV101 have similar fold as other known TSPs. The structures are coloured after the β-score. The beta-helices, hence the receptor binding domain of the TSPs had a higher prediction score compared to the head-binding domains.

### Figure S4

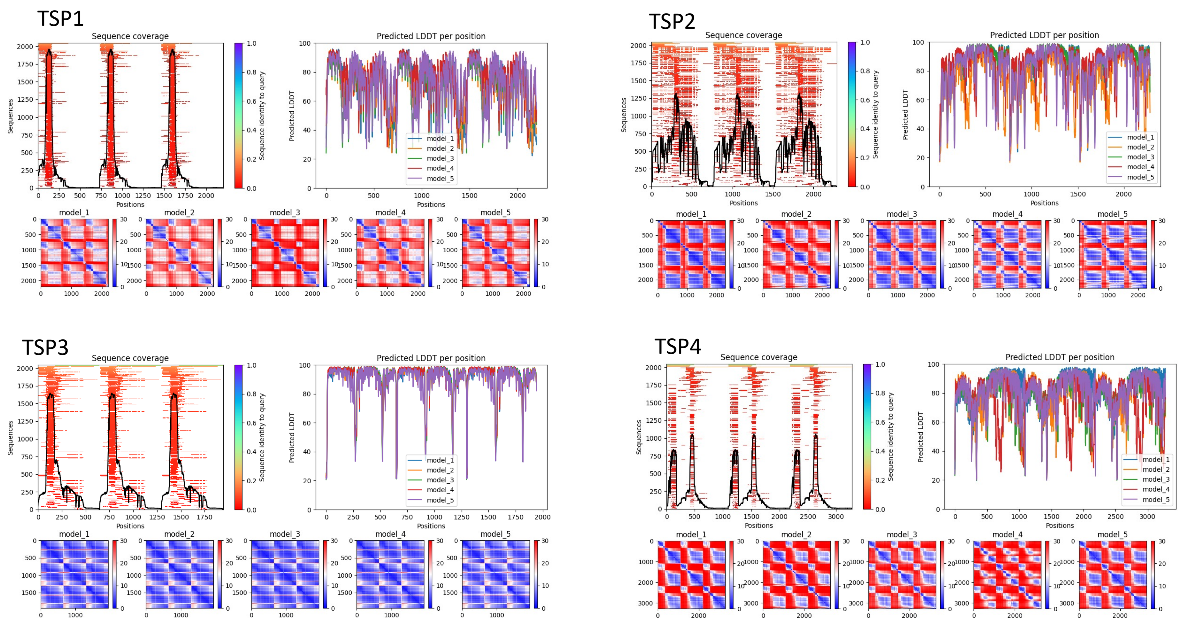


**Figure S4: Prediction quality metrics plots from Alphafold2 for the four TSPs.** The top-left graph represents the MSA coverage for each of the sequences. In the top-right is the pLDDT per position plot for the five final models for each TSP1.In the bottom is the Predicted Aligned Error matrix for the five final models for the four TSP1.
